# The TRIM32 granule promotes TDP-43 inclusion formation and amyloid aggregation through shuttle condensates

**DOI:** 10.64898/2026.02.11.705390

**Authors:** Fei Meng, Ziyan He, Jiechao Zhou, Ximing Ye, Rongzhen Zhang, David W. Nauen, Juan C. Troncoso, Paul F. Worley, Wenchi Zhang

## Abstract

Aberrant protein aggregation and inclusion formation are hallmarks of amyotrophic lateral sclerosis (ALS) and frontotemporal lobar degeneration (FTLD), which share overlapping genetic and pathological features. Similar aggregates are increasingly recognized in Alzheimer’s disease (AD) and limbic-predominant age-related TDP-43 encephalopathy (LATE). However, it remains unclear whether a shared molecular pathway drives this pathological aggregation. Here, we report the formation of stress-responsive cytoplasmic TRIM32 granules, distinct from stress granules. The E3 ubiquitin ligase TRIM32, together with the shuttle factor UBQLN2 and the autophagy adaptor p62/SQSTM1, drive condensation of TRIM32 granules that depend on E3 ligase activity and a network of intermolecular interactions. These condensates act as scaffolds that capture client proteins, including TDP-43 and ANXA11, and modulate their mobility. A unique hydrophobic loop within TRIM32’s substrate-binding domain mimics low-complexity motifs in ANXA11 and TDP-43, enabling selective retention via competitive binding mediated by the UBQLN2 STI1 domain. Moreover, TRIM32 granules promote amyloid aggregation of TDP-43, an effect that is exacerbated by a pathogenic UBQLN2 mutation. In brains from individuals with diverse neurodegenerative diseases, TRIM32 co-localizes with pathological phospho-TDP-43 (pTDP-43) inclusions, supporting a model that TRIM32-driven granules function as selective proteostasis sorting compartments that broadly contribute to TDP-43 proteinopathy.

## Introduction

Aberrant accumulation of aggregated proteins is a hallmark of many neurodegenerative disorders, including amyotrophic lateral sclerosis (ALS) and frontotemporal lobar degeneration (FTLD), which share overlapping clinical and pathological characteristics and are now recognized as part of a disease continuum^1^. Cross-sectional studies performed over the last decade estimate that up to 50% of ALS patients develop cognitive impairment associated with FTLD. Similarly, up to 30% of FTLD patients develop motor dysfunction^2^. Considerable progress has been made in unraveling the genetics of ALS/FTLD associated with causative mutations and risk variants in a wide range of genes, implicating autophagy and RNA processing as the central pathways^3^. Within the ALS–FTLD spectrum, pathological mislocalization of TAR DNA-binding protein 43 (TDP-43) from the nucleus to the cytoplasm, along with its accumulation in ubiquitin-positive inclusions, is observed in over 95% of ALS cases and approximately 45% of FTLD cases, as well as a number of other neurodegenerative diseases including Alzheimer’s disease (AD) and limbic-predominant age-related TDP-43 encephalopathy (LATE)^1, 4, 5^. Furthermore mutations in TDP-43 cause familial forms of FTLD and ALS, supporting a central role of TDP-43 in disease pathogenesis^6^.However, little is known about whether there exists a shared molecular pathway that drives pathological aggregation of TDP-43.

TDP-43 is a ubiquitous RNA-binding protein composed of two RNA recognition motifs (RRMs) that bind UG-repeat RNA in a sequence-specific fashion and are indispensable for its essential roles in RNA processing including the repression of non-conserved cryptic exons^7^. TDP-43 is predominantly found in the nucleus but also undergoes nucleocytoplasmic shuttling and can be found in cytoplasmic and neuritic RNP granules. TDP-43 harbors a C-terminal low-complexity domain (LCD) that is highly aggregation-prone^8, 9^, mediating phase separation, and has been proposed to act as a reaction center for the formation of pathological aggregates^10, 11^. Previous studies have shown that TDP-43 can be induced to form fibrillar aggregates under various non-physiological conditions like high protein concentration or long incubation with shaking or with truncated constructs of TDP-43 protein^8, 9, 12^. However, the structures of these experimental TDP-43 fibrils are distinct from the fibrils that have been extracted from brains of individuals with FTLD^13, 14^. Accordingly, it remains unknown what causes disease-relevant TDP-43 aggregation. Remarkably, in ALS/FTLD caused by penetrant dominant genetic mutations, brain and/or spinal tissue often contains aggregates of both TDP 43 and mutant gene encoded proteins, including ANXA11 and UBQLN2^15–18^.

ANXA11 is a member of the annexin family of proteins that share the biochemical property of Ca^2+^ dependent binding to membranes. ANXA11 harbors a unique N-terminus that contains a low-complexity domain. At its C-terminus, ANXA11 contains a structured annexin repeat domain that binds to negatively charged phosphatidylinositol 3-phosphate in the lysosomal membrane in a calcium-dependent manner^19,20^. ANXA11 shares common biophysical features with TDP-43, including their ability to undergo liquid– liquid phase separation via relatively weak but multi-valent interactions mediated by their low-complexity domains^21^. Moreover, TDP-43 can be found within axonal RNP transport granules where ANXA11 tethers these RNP granules via a low complexity N-terminal domain to lysosomes for long distance trafficking and delivery to distal locations within neurons^19^. Pathogenic variants of ANXA11 were first discovered in familial or sporadic ALS cases^22, 23^ with ANXA11 aggregates, with or without co-localization with TDP-43 aggregates^17, 22, 24, 25^. Recent studies demonstrated that TDP-43 and ANXA11 are similarly abundant and colocalized in inclusions of FTLD-TDP subtype C^26, 27^ and FTLD-motor neuron disease primary lateral sclerosis (FTLD-PLS)^28^. Remarkably, structural examination of TDP type C amyloid filaments by cryo-electron microscopy (cryo-EM) revealed a heteromeric filament structure consisting of co-assembled TDP-43 and ANXA11 fragments^29^, suggesting that their pathologies may share upstream aggregation mechanisms.

Dominant mutations in UBQLN2 cause X-linked ALS/(FTLD)^15, 30^, characterized neuropathologically by the deposition of TDP 43 and UBQLN2 aggregates in several CNS regions. UBQLN2 is a member of the ubiquitin-like protein family (Ubiquilins), featuring an amino-terminal ubiquitin-like (UBL) domain and a carboxyl-terminal ubiquitin-associated (UBA) domain, which mediate binding to proteasomes and ubiquitinated substrates, respectively. The middle region of UBQLN2 contains two STI1 domains^15, 31^, which form a hydrophobic groove that engages hydrophobic regions of client proteins^31, 32^. Fittingly, UBQLN2 functions as a shuttle factor, facilitating the delivery of misfolded protein cargos from stress granules to either the proteasome or the autophagy-lysosome system for degradation^33–35^. In addition, UBQLN2 harbors a unique PXX domain comprising 12 proline-rich tandem repeats between STI1 domain and UBA domain where most ALS/FLTD-associated pathological mutations reside^15, 36^. Interestingly, UBQLN2 inclusions are a near universal feature in TDP-43-positive ALS, as well as ALS /FTLD linkedto C9orf72 expansions (C9-ALS) ^15, 30, 36–38^, and also co-localize with other ALS/FTLD-linked proteins, such as TDP-43, p62 or ubiquitin^15, 30, 36–38^. Thus, while UBQLN2 appears to be involved in ALS/FTLD pathogenesis regardless of etiology, it is still unclear whether there is a unifying pathogenic mechanism associated with UBQLN2 mutations^39^, and how ALS/FTD-associated UBQLN2 mutations contribute to TDP-43 proteinopathy^40^.

Tripartite motif-containing 32 (TRIM32) is an E3 ubiquitin ligase belonging to the TRIM-NHL family, characterized by the presence of a substrate recognition region NHL domain at a variable C-terminal region^41^. Mutations in TRIM32 are causative of limb-girdle muscular dystrophy type 2H (LGMD2H)^42–44^ and Bardet-Biedl syndrome type 11^45^, with neurological phenotypes indicating its role in neuroprotection. In neurons, TRIM32 deficiency results in impaired dendritic arborization and synaptic plasticity^46–49^. TRIM32 promotes ubiquitination of ULK1 and ATG7 to facilitate autophagy^50, 51^. In addition, impaired TRIM32-mediated regulation of p62/SQSTM1 activity has been suggested as a pathological mechanism of disease-associated variants^52^. Notably, TRIM32 and p62 levels are elevated in a UBQLN2-mediated FTLD model^53^, which suggests TRIM32’s involvement in neurodegenerative mechanisms tied to impaired proteostasis and autophagic clearance.

Here, we investigate how TRIM32 contributes to pathological aggregation of ALS/FTLD-linked proteins and define a phase-separated condensation of TRIM32 granules assembled on scaffold-client organization under oxidative and hyperexcitability stresses. TRIM32 forms mixed polyubiquitin chains on multiple autoubiquitination sites and, together with UBQLN2 or p62/SQSTM1, undergoes E3 ligase activity–dependent condensation of TRIM32 granules. This behavior is driven by multivalent interactions mediated by UBA domains. We identify a distinctive hydrophobic loop within TRIM32’s substrate-binding domain that mimics low-complexity motifs in the client proteins ANXA11 and TDP-43 and promotes their selective retention within TRIM32 condensates through intermolecular binding. UBQLN2 further tunes condensate dynamics via STI1 domain-mediated competitive binding, producing differential mobility among condensate components. These properties define a new class of dynamic assemblies that we term “shuttle condensates.” In contrast to wild-type UBQLN2, pathogenic UBQLN2 variants show reduced mobility within TRIM32 condensates. Critically, TRIM32 condensates with UBQLN2 and p62 unexpectedly promotes amyloid aggregation of TDP-43 as an emergent property, an effect enhanced by pathogenic UBQLN2 variants and partially rescued by mutation of the TRIM32 mimic loop. In human brain tissue with TDP-43 proteinopathy, TRIM32 and UBQLN2 co-localize with rounded pathologically phosphorylated TDP-43 inclusions. Colocalizing inclusions are prominent in the amygdala, an early-affected region in AD, and in younger cases of LATE and FTD, underscoring a direct link between TRIM32 granules, inclusion formation, and pathological aggregation. Together, these findings support a model in which TRIM32 granules act as selective sorting compartments for misfolded protein that regulate client fluidity and facilitate the accumulation of insoluble aggregates, providing a generalizable framework for TDP-43 proteinopathies.

## Results

### Endogenous neuron TRIM32 granules induced by oxidative and hyperexcitability stress

To establish a physiological basis for examining the association of TRIM32 with UBQLN2, we immunoprecipitated endogenous UBQLN2 from mouse brain lysate. TRIM32 was recovered with UBQLN2, supporting their association in brain (Fig. 1A). We next asked if endogenous TRIM32 can form condensates, although it has been reported that TRIM32 formed cytoplasmic puncta when overexpressed in cultured cells^54, 55^. After validating the TRIM32 antibody (Extended Data Fig. 1A), we exposed primary mouse cortical neurons to oxidative stress and neuronal activity stimulus by treating with arsenite and bicuculline, respectively, which require cell to initiate quality control system for protein overturn and misfolding. Under basal conditions or weak stimulus, TRIM32 and UBQLN2 remained diffusely distributed throughout the cytoplasm with no significant puncta formation (Fig. 1B, 1C). However, under stronger combined treatment, endogenous TRIM32 formed discrete cytoplasmic puncta that colocalized with UBQLN2 (Fig. 1B, 1C), named as TRIM32 granules. The percentage of neurons with TRIM32 granules and the size of TRIM32 granules was significantly increased in a stress-dependent manner(Fig. 1D, 1E), indicating that stress magnitude and duration influence TRIM32 condensate appearance and size. To further characterize these TRIM32 granules, we examined their relationship with canonical stress granules marked by G3BP1. Under higher stress conditions (200 μM arsenite 4h + 40 μM bicuculline 24h), TRIM32 granules appeared as distinct cytoplasmic puncta that did not colocalize with G3BP1-and TDP-43-positive stress granules(Fig. 1D), suggesting that TRIM32 condensates with UBQLN2 as a constitutive component represent a unique granule population separate from canonical stress granules.

**Figure 1.**
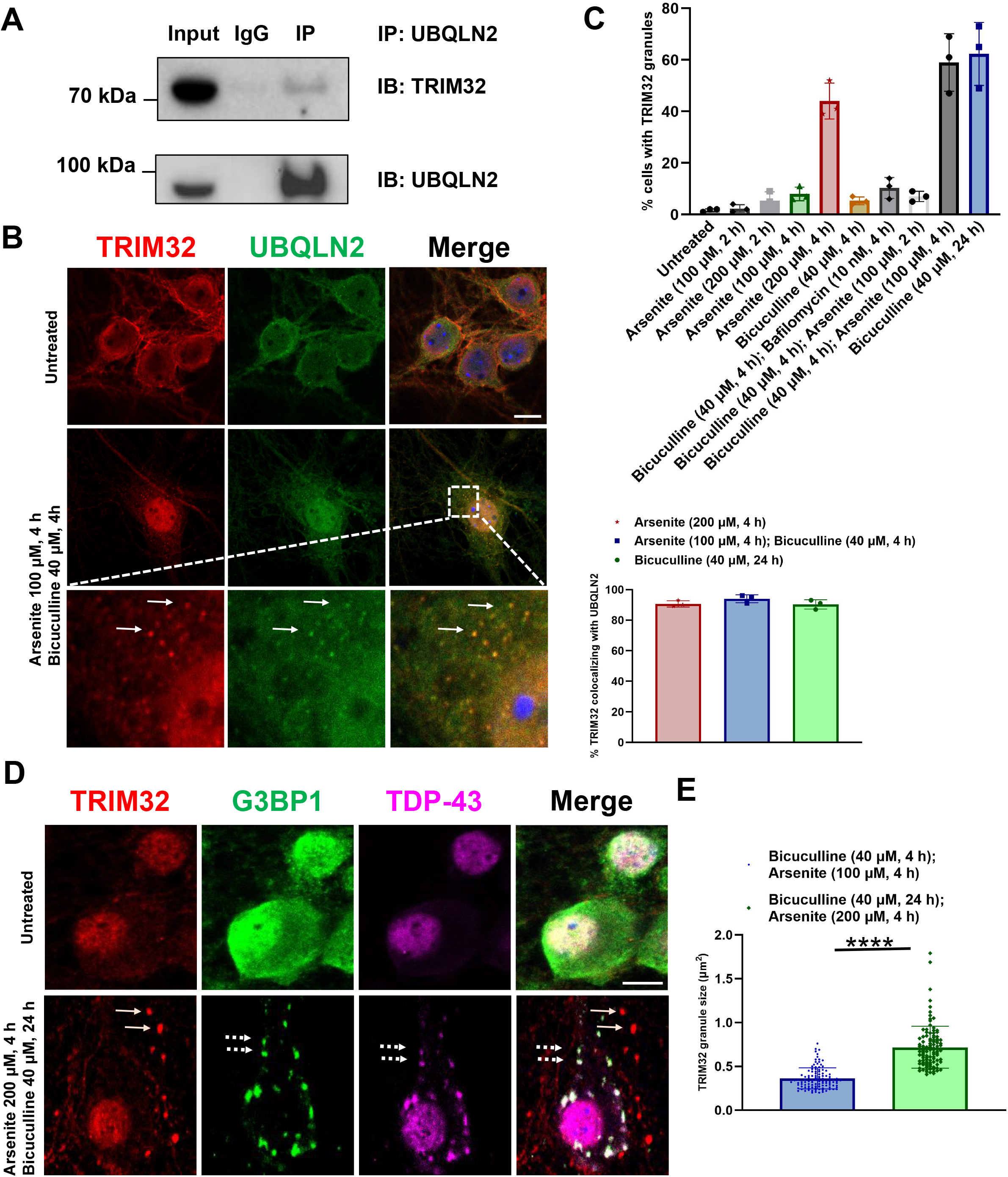
Characterization of neuron TRIM32 granules formed under oxidative stress and hyperactivity. **A)** Co-IP of TRIM32 and UBQLN2 from 9-month-old mouse brain lysate using UBQLN2 antibody. **(B)** Representative images of TRIM32 (red), UBQLN2 (green) and TDP-43 (purple) in primary mouse cortex neuron culture in untreated and treatment of 100 μM Arsenite 4 h + 40 μM Bicuculline 4 h. White arrows indicate cytoplasmic TRIM32 granules colocalizing with UBQLN2; Nuclei are stained in blue (DAPI) in all merged images. Box indicates enlarged area in inset. Bar, 10 μm. **(C)** Quantitative analysis of the number of neuron cells (%) with TRIM32 granules and the percentage of TRIM32 granules colocalizing with UBQLN2 in condition of various Arsenite and Bicuculline treatment. Data are presented as mean ± s.e.m. p values were determined by one-way analysis of variance (ANOVA). p < 0.05 (*), p < 0.01 (**), p < 0.001 (***), p < 0.0001 (****); ns, not significant; at least 60 cells/condition were counted for each replicate. Shown are combined data from three independent experiments. **(D)** Representative images of TRIM32 (red), G3BP1(green) and TDP-43 (purple) in primary mouse cortex neuron culture in 200 μM Arsenite 4 h and 40 μM Bicuculline 24 h exposure. White arrows indicate cytoplasmic TRIM32 granules; White dash arrows indicate stress granules; Nuclei are stained in blue (DAPI) in all merged images. Bar, 10 μm. **(E)** Quantitative analysis of the mean area of TRIM32 granules between treatment of 100 μM Arsenite 4 h + 40 μM Bicuculline 4h and 200uM Arsenite 4 h + 40 μM Bicuculline 24 h from three biological replicates. Data are presented as mean ± s.e.m. p values were determined by two-tailed unpaired Student’s *t* test. P < 0.05 (*), p < 0.01 (**), p < 0.001 (***), p < 0.0001 (****); ns, not significant; at least 100 granules were counted for each condition. Shown are combined data from three independent experiments.

### TRIM32 assembles mixed linkage polyubiquitin chains

TRIM32 E3 ligase (Fig. 2A) possesses a RING domain that can recognize the ubiquitin loaded E2 conjugating enzyme activated by the ubiquitin E1 enzyme UBE1 in an ATP-dependent process, and allows for the direct transfer of ubiquitin to substrates or an acceptor ubiquitin to form polyubiquitin (polyUb) chains^41, 56^. The ubiquitin ligase activity of TRIM32 has been studied in cell-based assays across a range of substrates, revealing variable outcomes regarding the polyubiquitin chain specificity^57^. While some studies indicate TRIM32 catalyzes K48 ubiquitin chain formation and targets proteins for proteosomal degradation^58, 59^, other reports suggest that TRIM32 mediates K63 or K27/K63 Ub chains and the ensuing lysosomal degradation or signaling processes^50, 51, 60–63^.

**Figure 2.**
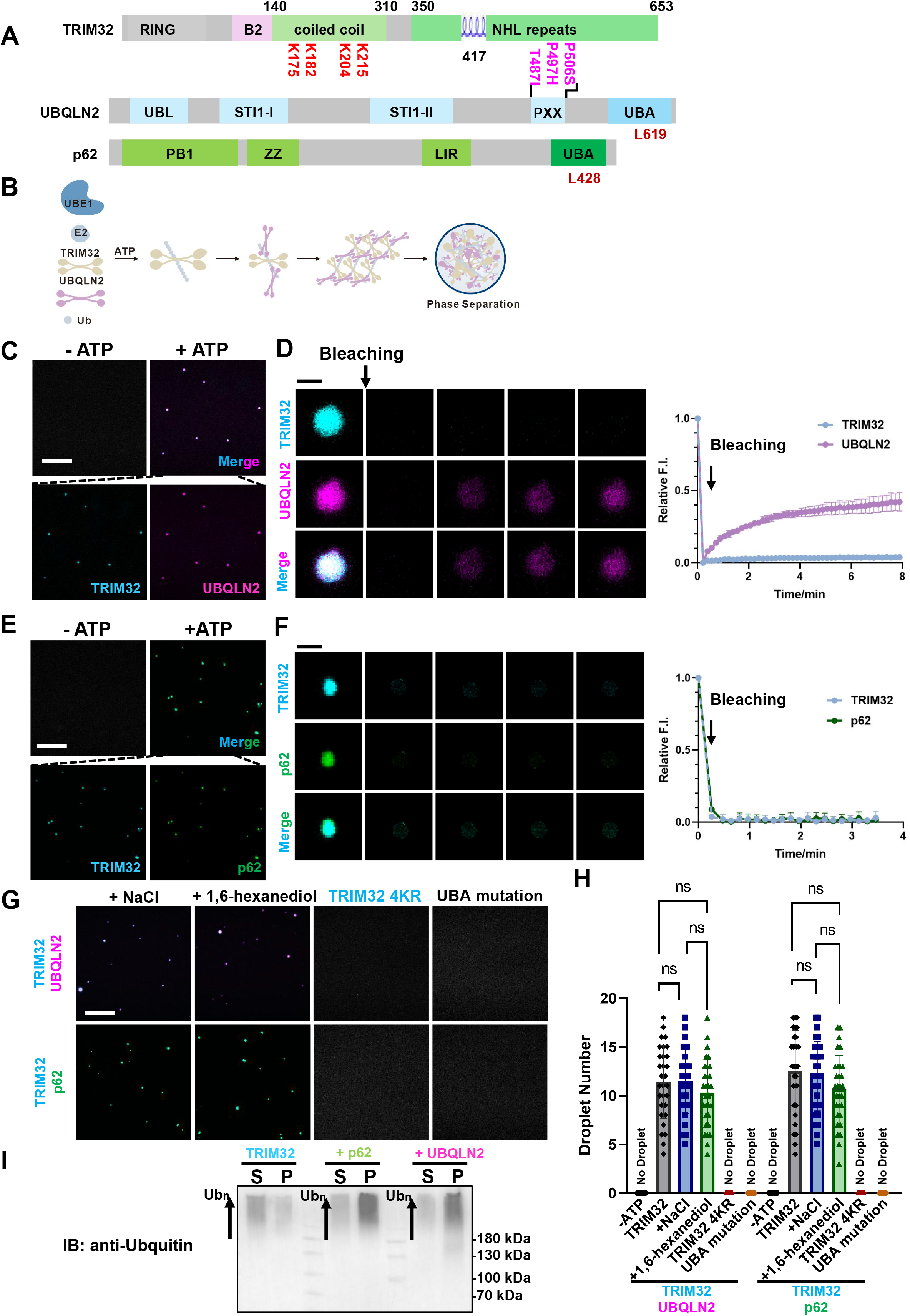
TRIM32 undergoes ligase activity dependent phase separation. **(A)** Schematic of human TRIM32, UBQLN2 and p62 highlighting domain architecture. The four lysine residues in the TRIM32 coiled-coil domain identified by mass spectrum analysis as autoubiquitination sites in the current study are indicated in red. The hydrophobic helix loop in TRIM32 NHL domain is labeled in blue. UBQLN2 Point mutations associated with human ALS/FTD pathogenesis are highlighted in purple. The conservative residues L619 at UBQLN2 UBA domain and L428 at p62 UBA domain mediating the interaction with ubiquitin are indicated in red. **(B)** Schematic illustration of TRIM32 activity induced phase separation mediated by multivalent interaction of UBA domain with auto-polyubiquitin chain on TRIM32. **(C)** TRIM32 ligase activity-dependent formation of phase separation in *in vitro* TRIM32 (cyan) autoubiquitination reaction with the addition of UBQLN2 (purple). Scale bar, 20 μm. **(D)** Left: fluorescence intensity recovery of a TRIM32-UBQLN2 condensate after photobleaching. Right: Quantification of fluorescence intensity recovery of photobleached TRIM32 and UBQLN2. *n* = 3. Data are presented as mean ± s.e.m. Scale bar, 2 μm. **(E)** TRIM32 ligase activity-dependent formation of phase separation in *in vitro* TRIM32 (cyan) autoubiquitination reaction with the addition of p62 (green). Scale bar, 20 μm. **(F)** Left: fluorescence intensity recovery of a TRIM32-p62 condensate after photobleaching. Right: Quantification of fluorescence intensity recovery of photobleached TRIM32 and p62. *n* = 3. Data are presented as mean ± s.e.m. Scale bar, 2 μm. **(G)** Fluorescence microscopy images showing buffers containing additional 100 mM NaCl or 3% 1,6-hexanediol have no effect on phase separation formation. But mutations in UBQLN2 or p62 UBA domain, and the quadruple point mutation TRIM32 4KR (K175R/K182R/K204R/K215R) inhibit phase separation formation. Scale bar, 20 μm. **(H)** The number of TRIM32 condensates showed in (**B), (D), (F)** was quantified. Each spot represents the number of condensates in one imaged area across 10 fields. Data are representative of three independent experiments as mean ± s.e.m. p values were determined by one-way analysis of variance (ANOVA). p < 0.05 (*), p < 0.01 (**), p < 0.001 (***), p < 0.0001 (****); ns, not significant. **(I)** Droplets sedimentation assay showing that pellet and supernatant of TRIM32 condensates formed during autoubiquitination reaction in presence of UBQLN2 or p62 were analyzed by western blot using anti-ubiquitin, indicating polyubiquitin (Ubn) enriched in pellets. S, supernatant; P, pellet.

To clarify the polyubiquitin chain linkage specificity of TRIM32 in a defined system, we expressed full-length recombinant TRIM32, UBE1, the E2 enzyme UBE2D3 selected based on cell-based assays^50, 51, 60–63^, and ubiquitin, and developed an *in vitro* reconstitution autoubiquitination assay (Extended Data Fig. 1B). TRIM32 exhibits high autoubiquitination ligase activity, exhausting ubiquitin monomers within 30 min and forms high molecular weight conjugates revealed as a smear of species at the top of the gel and loading well (Extended Data Fig. 1C, 1D). Mass spectrometry (MS) identified autoubiquitination sites on TRIM32 (Extended Data Fig. 1E) and mutation of these sites in TRIM32 4KR (K175R/K182R/K204R/K215R) prevents the autoubiquitination reaction (Extended Data Fig. 1C, 1D). MS also revealed mixed ubiquitin chains that contain K11, K48, and K63 linkages of near equal percentage (Extended Data Fig. 1F), suggesting lack of absolute preference for receptor ubiquitin placement during linkage formation of ubiquitin transfer^64^. Notably, mass spectrum identified concurrent GG-modification at adjacent lysines K6 and K11 (Extended Data Fig. 1G), indicating a presence of branched polyubiquitin. No ladder of polyubiquitin species was detected (Extended Data Fig. 1C) indicating TRIM32 preferentially assembles polyUb chains on substrates. These data establish an *in vitro* TRIM32 autoubiquitination assay and demonstrate TRIM32 assembles mixed polyubiquitin chains.

### UBQLN2 or p62 modulates TRIM32 E3 ligase activity

The high level of TRIM32 ligase activity suggested the possibility of a regulatory mechanism to restrict ligase activity. Our previous examination of the E3 ubiquitin ligase TRIAD3A demonstrated that an intramolecular interaction between its ubiquitin-binding domain and its autoubiquitin chains can inhibit ligase activity^65^, which prompted us to explore whether a similar regulatory mechanism might apply to TRIM32 by UBQLN2 and p62/SQSTM1, two ubiquitin-binding proteins that harbor UBA (ubiquitin-associated) domains (Fig. 2A).

To test the regulatory role of UBA domains in TRIM32 E3 ligase activity, we conducted in vitro autoubiquitination assays in which recombinant UBQLN2 or p62 was included in TRIM32 autoubiquitination reactions. Addition of either UBQLN2 or p62 significantly reduced TRIM32 ubiquitination activity, as revealed by the relative persistence of monomer ubiquitin, reduced polyubiquitin species at the top of the SDS gel over time, and reduced ubiquitin-linked proteins in western blots of reactions (Extended Data Fig. 2A, 2B, 2C). Despite reduced activity, TRIM32 polyubiquitin chain topology remained mixed (Extended Data Fig. 2D). To further assess the role of UBA domains, we applied AlphaFold2^66^ to generate complex structures of UBA domains with ubiquitin (Extended Data Fig. 2E) revealing conservative ubiquitin interaction surfaces that are consistent with previously characterized UBA-ubiquitin interfaces^67^. We further used our assay to determine that mutations L619A or L428A of UBA domains of UBQLN2 or p62, respectively, abolish their inhibitory action (Extended Data Fig. 2A, 2B). These results support a model in which UBQLN2 and p62 can serve as negative regulators of TRIM32 E3 ligase by binding to its autoubiquitin chains.

### TRIM32 undergoes E3 ligase dependent phase separation with UBA domain proteins

Network multivalent interactions can drive phase separation^68^. Since UBQLN2 or p62 form dimers or polymers, respectively, and TRIM32 can form tetramers^41^ and assemble complex polyubiquitin chains at multiple sites, we hypothesized that multivalent interactions of UBA domains and TRIM32 autoubiquitin chain may promote phase separation (Fig. 2B), in association with slowing of TRIM32 autoubiquitination.

To test this hypothesis, we reconstituted the autoubiquitination reaction using recombinant full-length TRIM32-mCerulean3, in the presence or absence of UBQLN2 conjugated with Alexa Fluor 647(UBQLN2-647) or p62-EGFP (Fig. 2B).In the absence of UBQLN2 and TRIM32, respectively, TRIM32 and UBQLN2 alone at concentration lower than 2uM did not undergo phase separation with ATP) (Extended Data Fig. 3A, 3B). Notably, in the control reaction without adenosine triphosphate (ATP) to initiate autoubiquitination, phase separation of TRIM32 and UBQLN2 or p62 did not occur at concentrations of 0.2∼1uM (Fig. 2C, 2E). Upon addition of ATP, we observed spontaneous phase separation without the need for artificial crowding agents (Fig. 2C, 2E). Importantly, mutations L619A or L428A in the UBA domains of UBQLN2 or p62, or 4KR (K175R/K182R/K204R/K215R) at TRIM32 autoubiquitination sites, impaired condensate formation (Fig. 2G, 2H), indicating that phase separation requires direct interactions between UBA domains of UBQLN2 or p62 and the autoubiquitin chains on TRIM32. We performed droplet sedimentation assays to confirm TRIM32-UBQLN2 or TRIM32-p62 condensates are enriched for polyubiquitin conjugates (Fig. 2I). Increasing NaCl concentration or adding 1,6-hexanediol did not impair condensate formation (Fig. 2G, 2H). This assembly mechanism of TRIM32 granules contrasts with phase-separation driven by high concentration and intrinsically disordered region of proteins, which typically depend on weaker electrostatic or hydrophobic interactions that can be disrupted by salt or hexanediol.

### Context-dependent exchange of UBQLN2 in TRIM32 condensates

Fluorescence recovery after photobleaching (FRAP) of TRIM32-UBQLN2 condensates revealed recovery for UBQLN2 indicating dynamic exchange with the surrounding solution (Fig. 2D). By contrast, TRIM32 in the same TRIM32-UBQLN2 condensates did not recover. This is consistent with the notion that polyubiquitinated TRIM32 functions as the hub scaffold for the condensate. FRAP analysis of TRIM32-p62 condensates revealed no recovery of TRIM32 or p62 (Fig. 2H) and addition of p62 to TRIM32 and UBQLN2 prevented FRAP of UBQLN2 (Extended Data Fig. 3C and 3D). Findings indicate context dependence of UBQLN2 fluidity in condensates where the presence of p62 may contribute to a more gel-like/solid-like state, consistent with prior reports that characterize p62-containing condensates as inherently less dynamic due to their ability to form stable polymers or fibrillar scaffolds^69, 70^ as a basis for confinement of cargo for autophagic processing.

**Figure 3.**
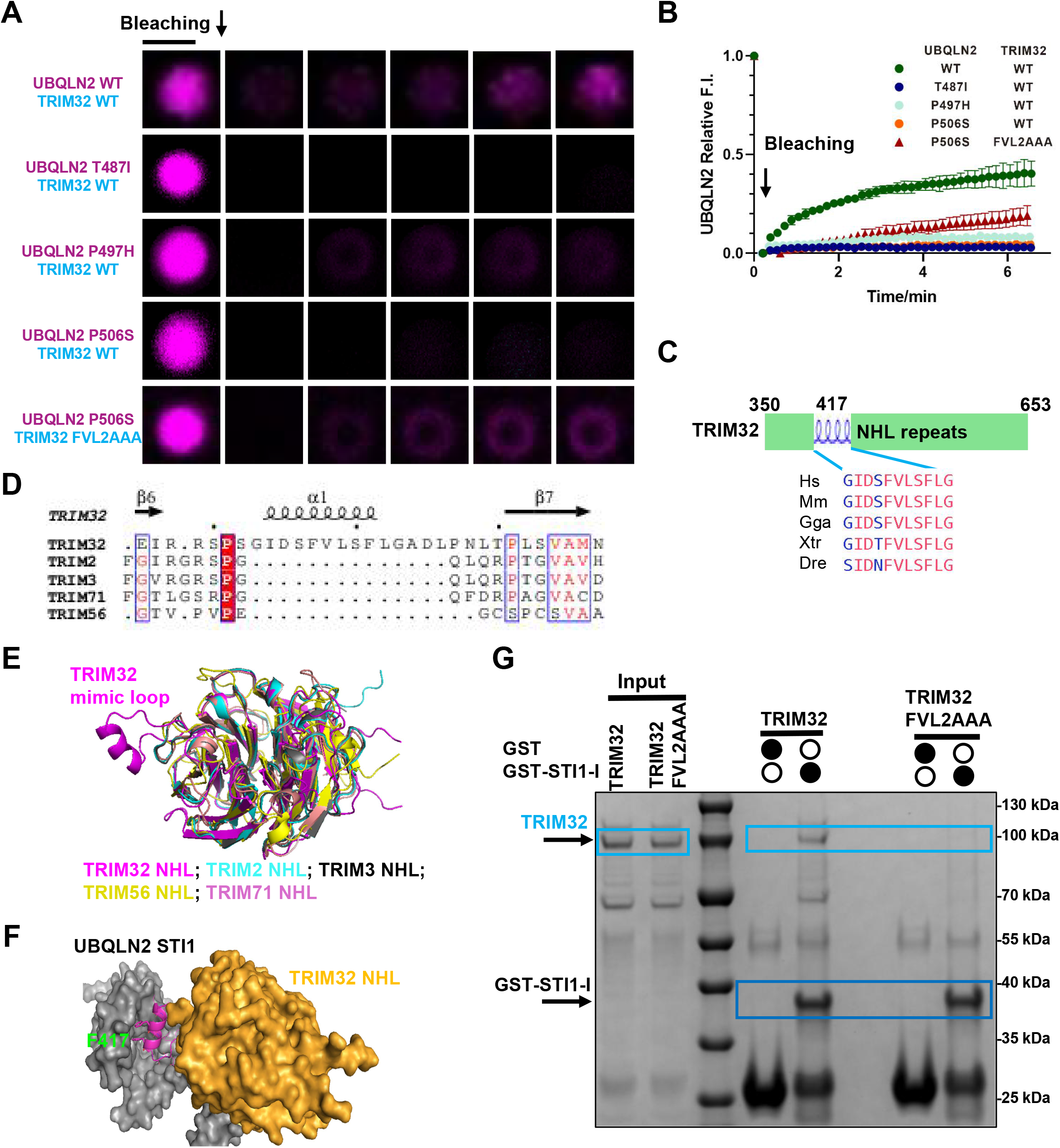
Reconstituted TRIM32 condensates underlie phase transition. **(A)** Fluorescence intensity recovery of WT UBQLN2 or UBQLN2 pathogenic mutants (purple) in a TRIM32-UBQLN2 condensate formed during TRIM32 autoubiquitination reaction in presence of UBQLN2 after photobleaching. Scale bar, 2 μm. **(B)** Quantification of fluorescence intensity recovery of photobleached WT UBQLN2 or UBQLN2 mutants. The mutation FVL2AAA (F417A/V418A/L419A) at the TRIM32 hydrophobic loop partly rescues the fluorescence recovery of UBQLN2 pathogenic mutant P506S. *n* = 3. Data are presented as mean ± s.e.m. **(C)** Position of the hydrophobic loop in TRIM32 NHL domain. The sequence alignment of this loop for typical species shows it is conservative. **(D)** Structure superposition of the NHL domains of human TRIM family members including TRIM32, TRIM2, TRIM3, TRIM56 and TRIM71, showing the unique hydrophobic helical loop only present in TRIM32. **(E)** The sequence alignment of partial NHL domains of human TRIM family members including TRIM32, TRIM2, TRIM3, TRIM56 and TRIM71, highlighting the unique hydrophobic helical loop only present in TRIM32. The sequence alignment of the whole NHL domains is in **Extended** figure 3F. **(F)** Complex structures of TRIM32 NHL domain and UBQLN2 STI1 domain generated by AlphaFold shows the hydrophobic loop in purple may form a-helix and occupy the hydrophobic binding pocket on UBQLN2 STI1-I domain. **(G)** Coomassie stained SDS-PAGE gels of in vitro GST pull-down demonstrating UBQLN2 STI1-I domain binds TRIM32, and mutation FVL2AAA at the hydrophobic helix of TRIM32 abolishes the interaction.

### UBQLN2 pathogenic mutations reduce exchange in TRIM32 condensates

To examine if pathogenic UBQLN2 mutations affect phase separation, we chose the T487I, P497H, and P506S mutations of the PXX domain (Fig 2A), which are common in UBQLN2 mutation-linked ALS/FTLD^15, 16, 38^. We added recombinant UBQLN2 mutants to TRIM32 *in vitro* autoubiquitination reconstitution reactions and found UBQLN2 mutants are sequestered within TRIM32 condensates like WT UBQLN2 (Extended Data Fig. 3E). However, in contrast to WT, UBQLN2 mutants did not recover after photobleaching (Fig. 3A, 3B). Examining this difference, we noted that mutations at the PXX domain are reported to disrupt intramolecular interactions with UBQLN2 STI1-I domain^32^, which we hypothesized might increase the availability of STI1-I domain to bind TRIM32. We identified an evolutionarily conservative loop in TRIM32 substrate binding NHL domain (Fig. 3C), that is unique to TRIM32 among TRIM family members containing the NHL domain as revealed by sequence alignment and predicted structure superposition (Fig. 3C, 3D, 3E and Extended Data Fig. 3F). Alphafold predicts this loop adopts a hydrophobic helix conformation that can occupy the binding pocket on UBQLN2 STI1-I domain (Fig. 3F). *In vitro* GST-pulldown experiments confirmed that TRIM32 binds UBQLN2 STI1-I domain, while the mutation TRIM32 FVL2AAA (F417A/V418A/L419A) at the TRIM32 hydrophobic loop reduced UBQLN2 binding (Fig. 3G). In the reconstitution assay, TRIM32 FVL2AAA partly rescued fluorescence recovery of UBQLN2 P506S (Fig. 3A, 3B), supporting a role for the TRIM32 NHL hydrophobic loop as a basis for intermolecular interactions within the condensate that impact UBQLN2 mobility.

### TRIM32 condensates coacerate UBQLN2 client proteins

Numerous proteins linked to ALS/FTLD pathogenesis co-aggregate with UBQLN2 including TDP-43 and ANXA11^17, 22, 24–27^. Emerging evidence reveals that the STI1-I domain of UBQLN2 optimally binds regions of moderately hydrophobic helicity^32^. ANXA11 and TDP-43 both contain α-helical regions in their low complexity domain (LCD) that are involved in self-association by helix-helix contact, and their conformation transition to beta-sheet promotes fibril assembly^71, 72^ (Fig. 4A). AlphaFold^66^ predicted their interaction with UBQLN2 STI1-I domain (Extended Data Fig. 4A) and *in vitro* GST pulldown experiments confirmed the physical interaction of UBQLN2 STI1-I domain with ANXA11 and TDP-43 (Extended Data Fig. 4B). Interestingly, TRIM32 hydrophobic loop and α-helical regions in LCD of TDP-43 or ANXA11 occupy the same site on UBQLN2 STI1-I domain (Fig. 3F, 4B, and Extended Data Fig. 4A). Consistent with a potential competition within this interaction network, co-immunoprecipitation showed reduced recovery of TDP-43 with UBQLN2 in the presence of TRIM32 (Fig. 4C).

To examine the phase behavior of TDP-43 and ANXA11 within TRIM32 condensates, we added mCherry-TDP-43 or ANXA11 conjugated with Alexa Fluor 546(ANXA11-546) to TRIM32-p62 *in vitro* reconstitution reaction with or without presence of WT UBQLN2 or UBQLN2 P506. We observed TDP-43 and ANXA11 coacervate into TRIM32-p62 condensates (Extended Data Fig. 4C, 4D). FRAP analysis of TDP-43 or ANXA11 revealed no fluorescence recovery in TRIM32-p62 condensates in absence of UBQLN2 or with UBQLN2 P506 mutant (Fig. 4D, 4E). By contrast, both TDP-43 and ANXA11 in TRIM32–UBQLN2-p62 condensates exhibited fluorescence recovery. Findings indicate that UBQLN2 promotes fluidity of ANXA11 and TDP-43 in the TRIM32-p62 condensates, consistent with UBQLN2-mediated competition for binding of client proteins to TRIM32. The role of competitive binding within the condensate is further supported by the observation that UBQLN2 does not display mobility without client proteins in TRIM32-p62 condensates (Extended Data Fig. 3C, 3D).

Collectively, these results suggest that the relative binding affinity of client proteins to the shuttle factor UBQLN2 versus the scaffold TRIM32 determines its fluidity and retention within the condensate. We refer the hierarchical organization of the fluidity of TRIM32-UBQLN2-p62 condensates as “shuttle condensates”, highlighting mobility heterogeneity of condensate components and the selective retention of clients based on their relative binding avidity to TRIM32 and UBQLN2.

### TRIM32-UBQLN2-p62 condensate promotes amyloid transition of TDP-43

To test whether TRIM32 condensates promote TDP-43 aggregation, we reconstituted TRIM32 condensates in the presence of TDP-43 and p62, together with either WT UBQLN2 or UBQLN2 (P506S). We then assayed amyloid protein formation by staining with Amylo-Glo, an amyloid aggregate indicator dye^103^. At 18 h after phase separation was induced by TRIM32 E3 ligase activity, a portion of TDP-43 containing TRIM32 condensates became Amylo-Glo-positive. Approximately 40% were positive with WT UBQLN2, whereas a larger fraction (∼60%) were positive with UBQLN2 (P506S) (Fig. 4F). Furthermore, we utilized immunogold labelling with TDP-43-specific antibodies, followed by negative staining electron microscopy, and observed fibrillar TDP-43 structures (Fig. 4H, 4I), indicating amyloid-like conversion of TDP-43. These data indicate that retention of TDP-43 within TRIM32 condensates promotes amyloid aggregation, and that this process is accelerated by a pathogenic UBQLN2 mutation.

### Cellular TRIM32 condensates parallel in vitro assays and recapitulate neuropathological features

To characterize the behavior of TRIM32 condensates in cells, we confirmed that UBQLN2 and UBQLN2 pathogenic mutants expressed alone in HEK293 cells localize diffusely within the cytoplasm (Extended Data Fig. 5A), consistent with prior findings^15, 73^. However, 24 h after co-transfection of TRIM32 with WT UBQLN2 the proteins exhibited colocalized spherical puncta (Fig. 5A). Previous studies have reported similar UBQLN2 puncta suggesting phase separation, but only in association with cell stress^15, 35^. Observations here suggest that TRIM32 is sufficient to drive phase separation of UBQLN2 in vivo without additional stress. Similar spherical puncta were evident in cells co-expressing TRIM32 and p62 (Extended Data Fig. 5B). Consistent with *in vitro* assays, mutation of the UBA domains of UBQNL2 or p62 to disrupt polyubiquitin binding reduced puncta formation (Extended Data Fig. 5C and 5D). Importantly, while WT UBQLN2 produced spherical puncta, UBQLN2 pathogenic mutations produced larger inclusions with irregular shape, suggesting a solid property of condensates (Fig. 5A). Bafilomycin A1 (BafA1) treatment to inhibit lysosome function did not significantly alter the morphology or size of UBQLN2 inclusions (Fig. 5B, 5C and Extended Data Fig. 5E), consistent with the notion that transgene expression of TRIM32 mimics and occludes the effect of lysosome stress to induce condensate formation.

**Figure 4.**
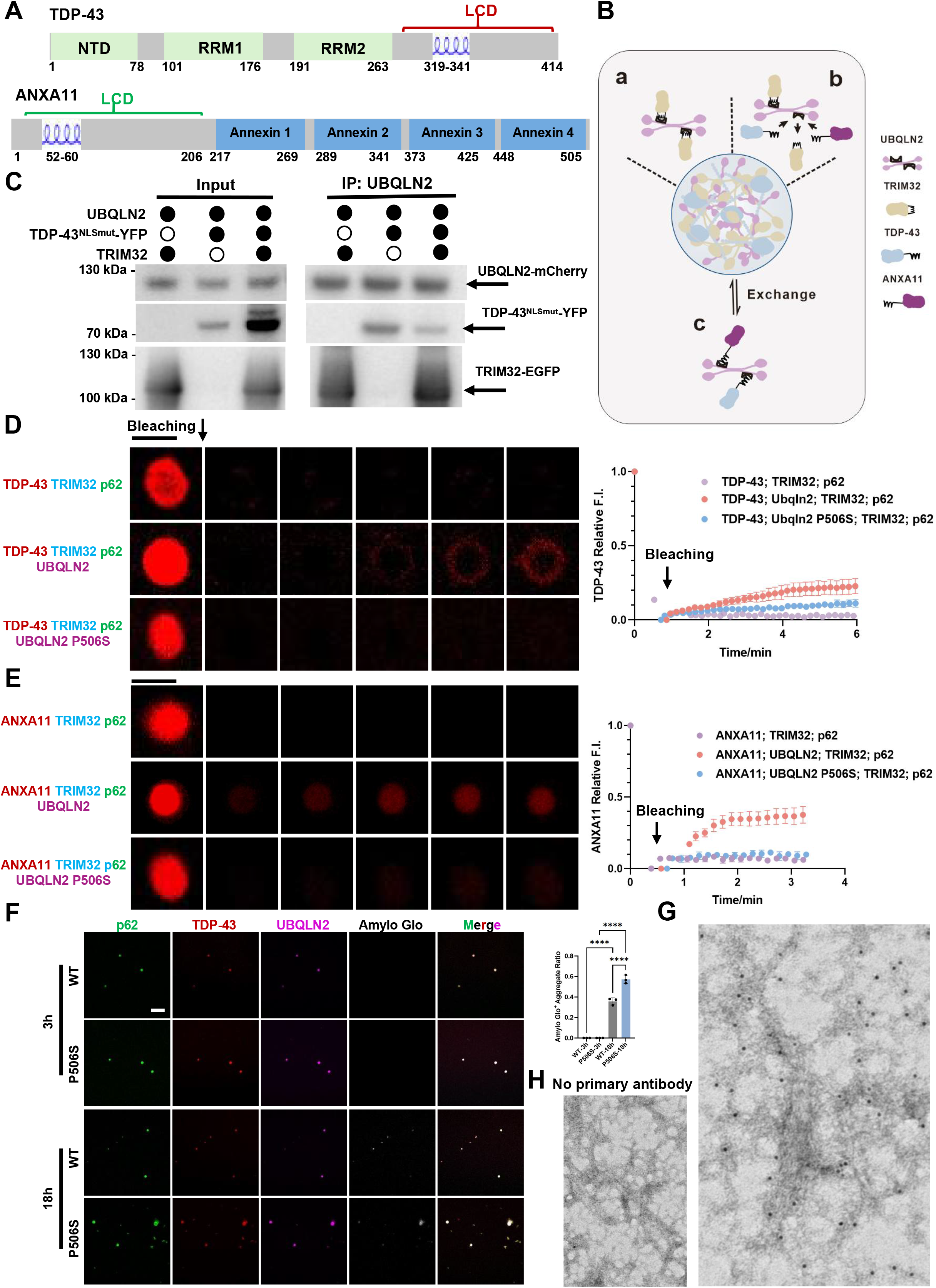
TRIM32 condensates modulate retention and aggregation of client proteins. **(A)** Position of the helix region in the low complexity domain (LCD) of the full-length ANXA11 or TDP-43. **(D)** Schematic representation of three modes of UBQLN2 binding governed by competitive interactions among the scaffold TRIM32, UBQLN2, and client proteins ANXA11 or TDP-43 within TRIM32–UBQLN2 condensates. (a) Both binding sites of the UBQLN2 dimer interact with TRIM32, anchoring UBQLN2 within condensates. (b) Client proteins ANXA11 or TDP-43 competitively occupy the same binding site on UBQLN2. (c) Both binding sites of the UBQLN2 dimer are occupied by ANXA11 or TDP-43, allowing UBQLN2–client complexes to diffuse into or out of condensates. **(E)** Representative western blotting images showing co-IP of UBQLN2 constructs co-expressed in HEK293 cells with TDP-43^NLSmut^-mCherry or TRIM32-EGFP, indicate competitive interaction of UBQLN2 with TDP-43 and TRIM32. **(F)** Left: fluorescence intensity recovery of ANXA11 (red) in the TRIM32-p62 or TRIM32-UBQLN2-p62 or TRIM32-UBQLN2 (P506S) -p62 condensate formed during the TRIM32 auto-ubiquitination reaction in vitro. Right: Quantification of fluorescence intensity recovery of photobleached ANXA11 in three kinds of condensates. *n* = 3. Data are presented as mean ± s.e.m. Bar=2 μm. **(G)** Left: Representative images of Amylo-Glo, an amyloid cross-β-sheet binding dye (white) staining at the indicated times, showing strong co-localization with TDP-43 (red) in TRIM32 condensates formed during the TRIM32 auto-ubiquitination reaction in vitro with addition of UBQLN2 (purple) and p62 (green). Right: Quantification of the ratio of Amylo-Glo positive TDP-43 condensates among UBQLN2-p62-TDP-43 colocalized condensates. Number of puncta are quantified from three independent experiments. Data are presented as mean ± s.e.m. p values were determined by one-way analysis of variance (ANOVA). p < 0.05 (*), p < 0.01 (**), p < 0.001 (***), p < 0.0001 (****); ns, not significant. **(H,I)** Representative TEM images showcasing immunogold labelling on TDP-43 fibrils extracted from TRIM32-UBQLN2-p62 condensates reconstituted in vitro.

**Figure 5.**
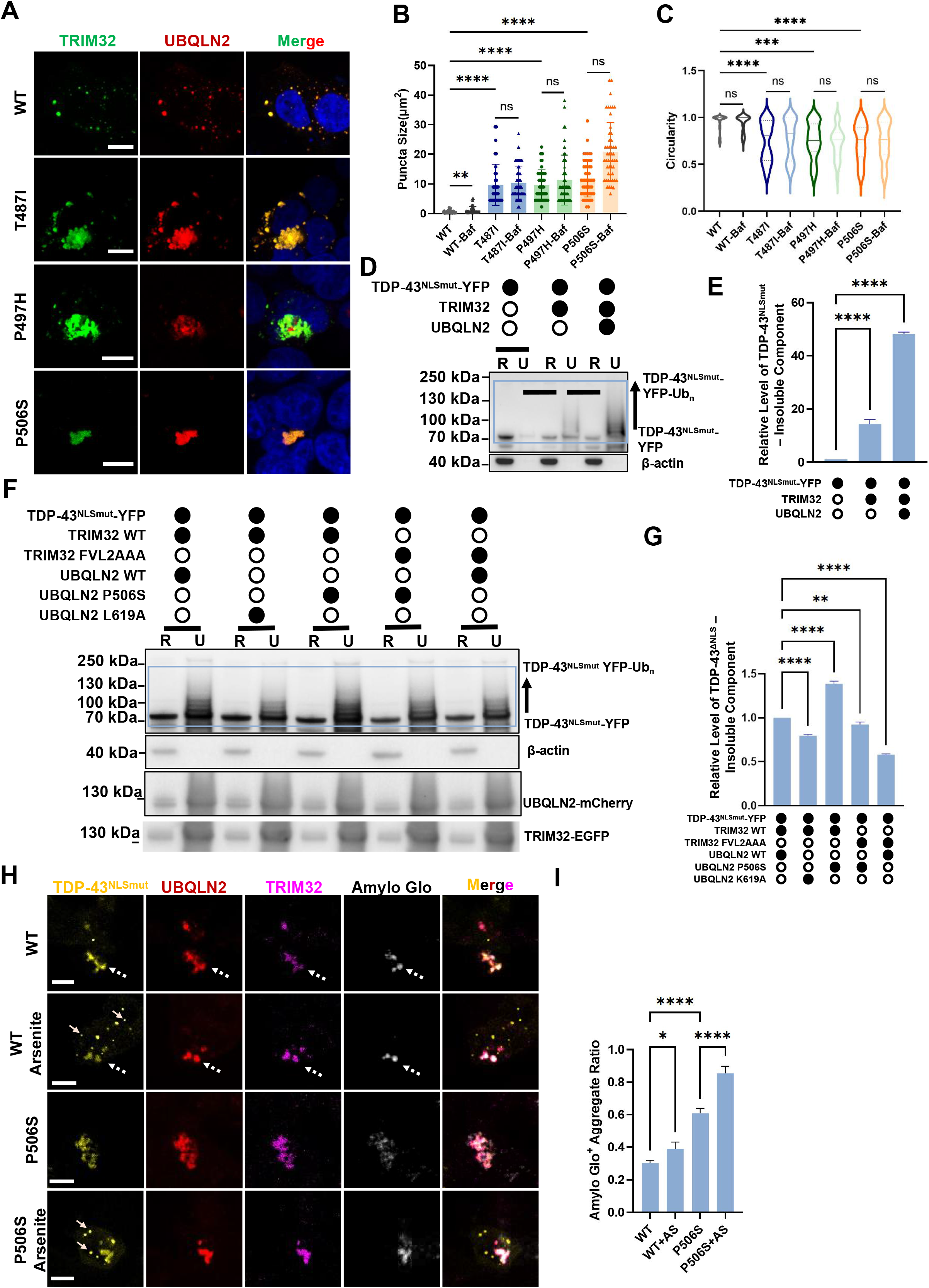
TRIM32 condensates recapitulate neuropathological features. **(A)** Representative immunofluorescence images of TRIM32 (green) and wt UBQLN2 or UBQLN2 pathogenic mutants (Red) with DAPI (blue) in HEK293T cells. Scale bar, 10 μm. **(B, C)** Quantification of TRIM32-UBQLN2 condensates with and without Bafilomycin A1 treatment (Figure A and Extended figure 5E). Data are presented as mean ± s.e.m. Statistical significance was determined using one-way ANOVA between mutation groups and WT group, followed by pairwise comparisons using unpaired two-tailed t-tests between Bafilomycin treated and untreated groups. p < 0.05 (*), p < 0.01 (**), p < 0.001 (***), p < 0.0001 (****); ns, not significant. Each spot represents one puncta. Shown are combined data from three independent experiments. **(D)** Representative western blot images to assess the solubility of TDP-43^NLSmut^ with or without cotransfection of TRIM32 or UBQLN2 in RIPA (R) and urea (u) buffers using anti-TDP-43 antibody. TDP43^NLSmut^-YFP-Ubn indicates ubiquitinated TDP-43^NLSmut^-YFP. **(E)** Quantification of relative levels of TDP-43^NLSmut^-YFP and ubiquitinated TDP-43^NLSmut^-YFP from three independent experiments showed in **(D)**. The box indicates the specific band for TDP-43^NLSmut^ and its ubiquitinated species for quantification. **(F)** Representative western blot images to assess the solubility of TDP-43^NLSmut^ with cotransfection of TRIM32 mutation FVL2AAA or UBQLN2 UBA point mutation L619A or pathogenic mutation P506S in RIPA (R) and urea (u) buffers, using anti-TDP-43 antibody. TDP-43^NLSmut^-YFP-Ubn indicates ubiquitinated TDP-43^NLSmut^-YFP. **(G)** Quantification of relative levels of TDP-43^NLSmut^-YFP and ubiquitinated TDP-43^NLSmut^-YFP from three independent experiments showed in **(F)**. The box indicates the specific band for TDP-43^NLSmut^-YFP and its ubiquitinated species for quantification. **(H)** Representative immunofluorescence images of HEK293T cells with cotransfection of TDP-43^NLSmut^ (yellow), TRIM32 (purple) and UBQLN2 WT or disease-associated mutant P506S (red) staining with with an amyloid aggregate indicator dye, Amylo-Glo (white) under no stress or arsenite stress (AS) condition. TDP-43 stress granules and TRIM32 condensates are indicated with white arrows and white dash arrows, respectively. Scale bar, 10 μm. **(I)** Quantification of the ratio of Amylo-Glo positive TDP-43 aggregates among TRIM32-UBQLN2-TDP-43 colocalized puncta showed in **(H)**. Number of puncta are quantified from three independent experiments. **(E,G,I)**Data are presented as mean ± s.e.m. p values were determined by one-way analysis of variance (ANOVA). p < 0.05 (*), p < 0.01 (**), p < 0.001 (***), p < 0.0001 (****); ns, not significant.

We next examined TRIM32-UBQLN2 condensates in HEK293 cells for inclusion of ANXA11 or TDP-43 using TDP-43 ^NLSmut^ , a mutant that localizes to cytoplasm^74^. We confirmed that either protein expressed alone in HEK293 cells is cytoplasmic and diffuse (Extended Data Fig. 5F, 5I), consistent with prior findings^11, 23^. However, when co-expressed with TRIM32 and UBQLN2, TDP-43 ^NLSmut^ or ANXA11 formed irregular inclusions that colocalize with TRIM32 and UBQLN2 (Extended Data Fig. 5F, 5G, 5I, 5J). To assess the physical state of cellular TDP-43^NLSmut^ and ANXA11, we performed detergent solubility assays^39, 75^. Results show that TDP-43 and ANXA11 are markedly enriched in RIPA insoluble fractions when coexpressed with TRIM32 and UBQLN2(Fig. 5D, 5E and Extended Data Fig. 5H). Pathogenic mutant UBQLN2 (P506S) and UBA domain mutant L619A increased or reduced the accumulation of insoluble TDP-43 aggregates, respectively, whereas the TRIM32 hydrophobic loop mutant (FVL2AAA) attenuated TDP-43 aggregate formation (Fig. 5F, 5G). These observations recapitulate *in vitro* studies and indicate that accumulation of UBQLN2 clients involves physical interactions within the TRIM32-UBQLN2 condensate and can induce transition to a detergent insoluble state.

Finally, we asked whether client proteins recruited to TRIM32-UBQLN2 condensates transition to an amyloid state in cells. Prior work indicates that oxidative stress induced by sodium arsenite can promote pathological TDP-43 aggregation. We co-expressed TRIM32 and TDP-43 ^NLSmut^ together with either WT UBQLN2 or UBQLN2(P506S), and stained cells with Amylo-Glo. Both WT UBQLN2 and UBQLN2(P506S) produced Amylo-Glo-positive TDP-43 aggregates that co-localized with TRIM32 and UBQLN2 (Fig. 5H, 5I). The Amylo-Glo-positive aggregate burden was higher in cells expressing UBQLN2 (P506S). Arsenite exposure further increased Amylo-Glo-positive aggregates with both WT UBQLN2 and UBQLN2 (P506S) (Fig. 5H, 5I). Arsenite also induced TDP-43 positive stress granules that were TRIM32/UBQLN2/Amylo-Glo-negative (Fig. 5H), consistent with endogenous observation that TRIM32 granules are distinct from stress granules (Fig. 1D), and supporting that amyloid-like TDP-43 accumulation occurs outside stress granules.

Collectively, these data support a model in which TRIM32-UBQLN2 condensates function as a sorting compartment. They permit exchange of soluble client proteins while preferentially retaining insoluble, pathological assemblies, thereby promoting aggregate accumulation under proteotoxic stress.

### p62 and pTDP-43 accumulation within TRIM32 condensates in neuron with autophagic flux inhibition

To assess the pathological relevance of TRIM32 granules, we returned to primary cortical neurons to determine whether the endogenous condensates identified in Fig. 1 associate with p62 and pathological pTDP-43. Under arsenite/bicuculline treatment, addition of bafilomycin A1 increased the fraction of TRIM32 puncta containing p62 (Fig. 7A,B). In parallel experiments, cytoplasmic pTDP-43 at Ser409/410 was rarely detected in untreated neurons, increased under the stress regimen, and was more frequent with bafilomycin A1. Representative pTDP-43 puncta colocalized with TRIM32 and UBQLN2 (Fig. 7C,D). Consistent with reconstitution experiments, these observations support that endogenous stress-responsive TRIM32 granules accumulate pathological client protein when autophagy-lysosomal pathway is inhibited and proteostatic burden increased.

**Figure 6.**
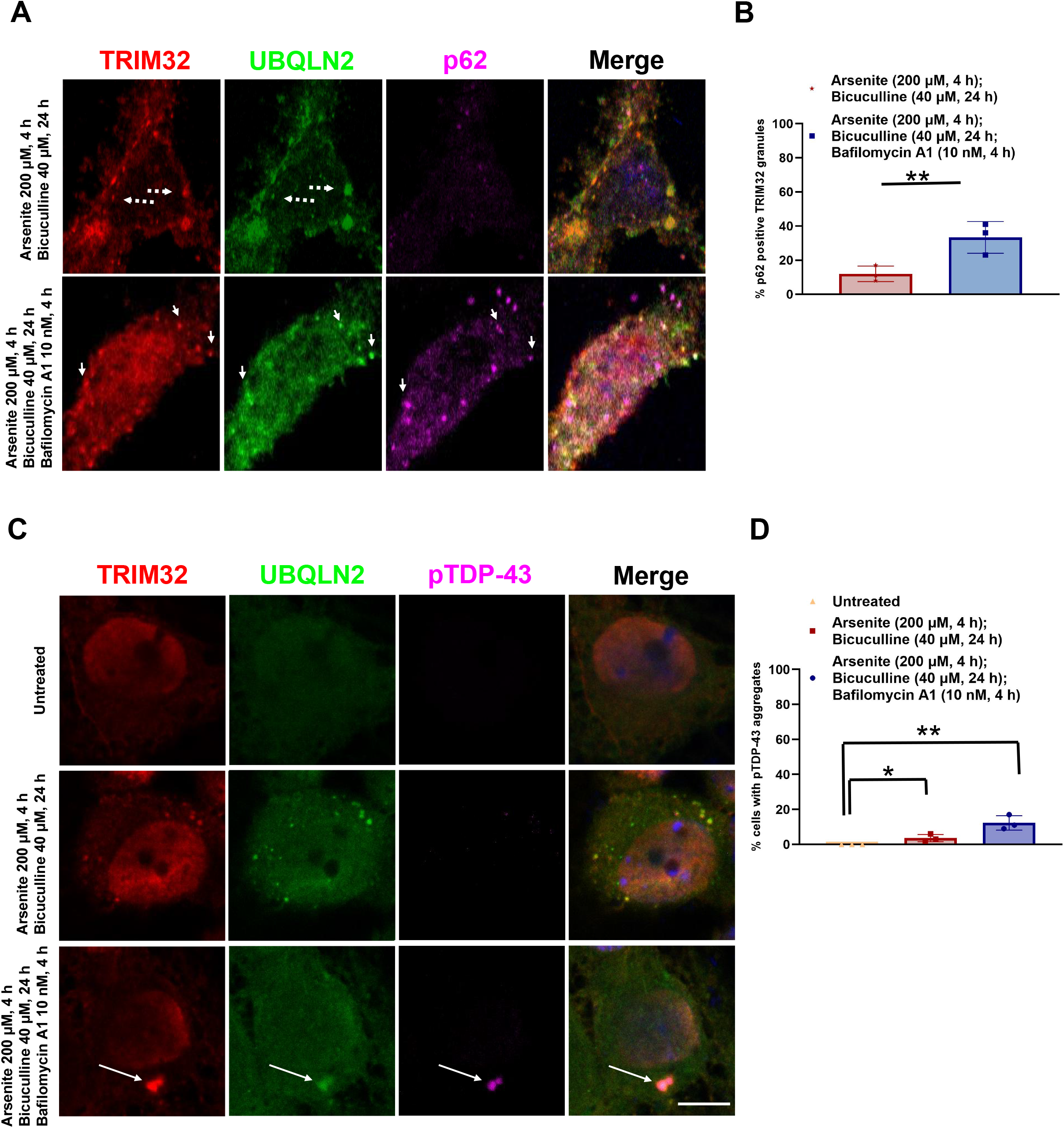
Endogenous p62 and pTDP-43 accumulation in primary cortex neuron. **(A)** Representative images of TRIM32 (red), UBQLN2 (green) and p62/SQSTM1 (purple) in primary mouse cortex neuron culture in 200 μM Arsenite 4 h, 40 μM Bicuculline 24 h and 10 nM Bafilomycin A1 4 h exposure. White solid arrows indicate cytoplasmic TRIM32 granules coloalizing with UBQLN2 and p62; White dash arrows indicate cytoplasmic TRIM32 granules coloalizing with UBQLN2; Nuclei are stained in blue (DAPI) in all merged images. Bar, 10 μm. **(B)** Quantitative analysis of the number of TRIM32 granules (%) with p62 in condition of Arsenite and Bicuculline with or without Bafilomycin A1 treatment shown in (E). Data are presented as mean ± s.e.m. p values were determined by one-way analysis of variance (ANOVA). p < 0.05 (*), p < 0.01 (**), p < 0.001 (***), p < 0.0001 (****); ns, not significant; at least 60 cells/condition were counted for each replicate. Shown are combined data from three independent experiments. (C) Representative images of TRIM32 (red), UBQLN2 (green) and pTDP-43 (purple) in primary mouse cortex neuron culture in 200 μM Arsenite 4h, 40 μM Bicuculline 24 h and 10 nM Bafilomycin A1 4 h exposure. White arrows indicate cytoplasmic TRIM32 granules coloalizing with UBQLN2 and pTDP-43; Nuclei are stained in blue (DAPI) in all merged images. Bar, 5 μm. **(D)** Quantitative analysis of the number of neuron cells (%) with cytoplasmic pTDP-43 aggregates in condition of Arsenite and Bicuculline with or without Bafilomycin A1 treatment shown in (J). Data are presented as mean ± s.e.m. p values were determined by one-way analysis of variance (ANOVA). p < 0.05 (*), p < 0.01 (**), p < 0.001 (***), p < 0.0001 (****); ns, not significant; at least 60 cells/condition were counted for each replicate. Shown are combined data from three independent experiments.

**Figure 7.**
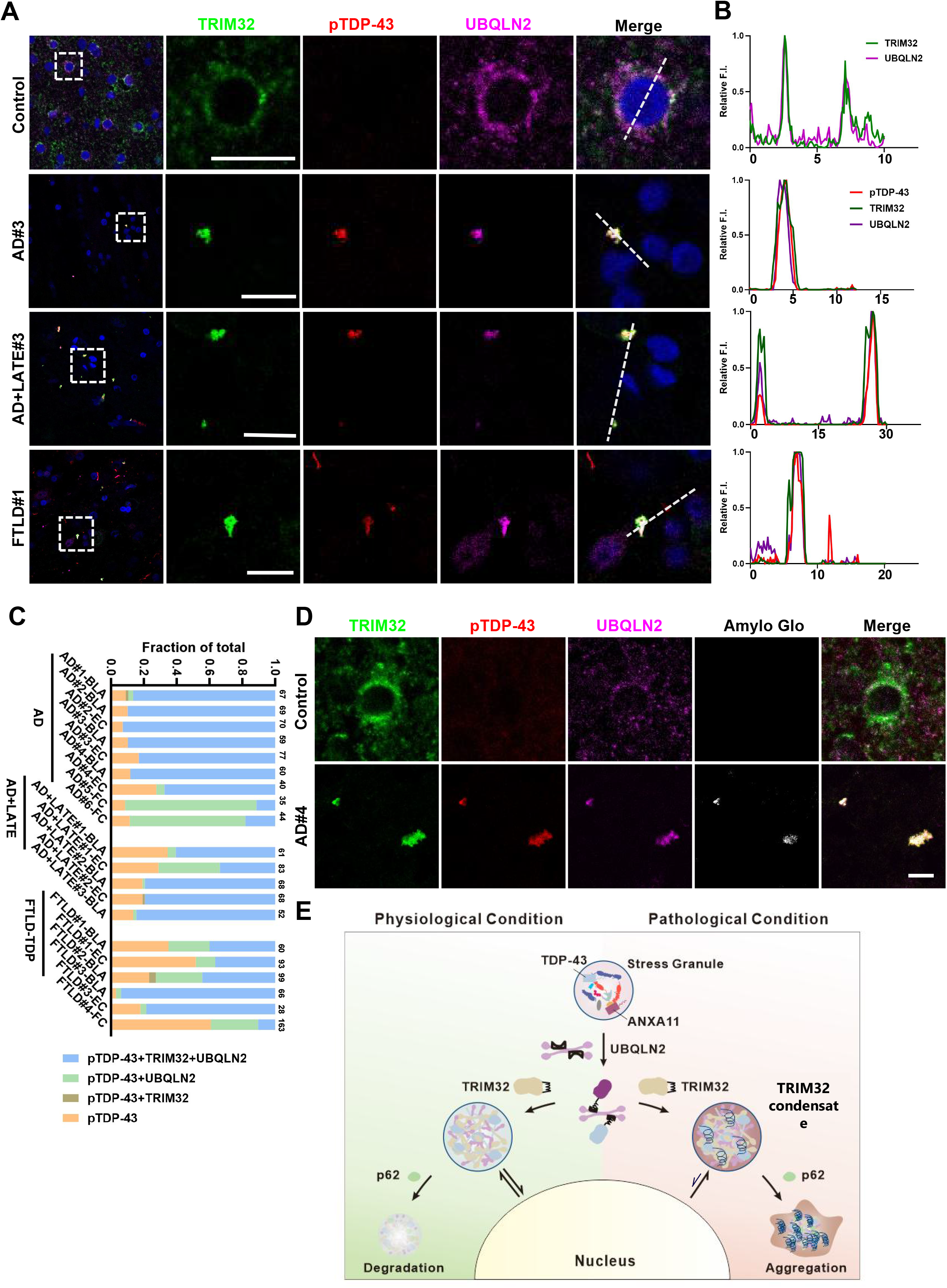
TRIM32 condensates occur in patients with TDP-43 proteinopathy. **(A)** Representative images showing TRIM32, UBQLN2 and pTDP-43(Ser409/410) within round-like pTDP-43 inclusions in the amygdala region of non-dementia (control), AD, AD with LATE, and FTD subjects with TDP-43 proteinopathy. Areas outlined by dotted white boxes are magnified to the right. Scale bar, 10 μm. **(B)** Radial distribution of the fluorescence intensities of TRIM32, UBQLN2 and pTDP-43(Ser409/410) indicated that TRIM32 co-localization with UBQLN2 and pTDP-43. **(C)** Quantification of the percentage of pTDP-43 positive aggregates co-localizing with/without TRIM32 or UBQLN2 in the basolateral and lateral nucleus (BLA), entorhinal cortex (EC) in the amygdala region, and frontal cortex (FC) from AD, AD with LATE, and FTD subjects with TDP-43 proteinopathy. Numbers above each column indicate the counts of pTDP-43 positive aggregates for the corresponding case. **(D)** Representative images of TRIM32, UBQLN2, pTDP-43(Ser409/410) and Amylo-Glo in the human amygdala region of non-dementia (control) and AD subjects. Scale bar, 5 μm **(E)** Schematic of the proteostasis mechanism mediated by TRIM32-UBQLN2-p62 shuttle condensates. The shuttle factor UBQLN2 with clients TDP43 and ANXA11 et al may exchange between various pools like stress granules, TRIM32 condensates and nuclear based on their relative binding avidity to scaffolds, which at last will be confined in p62 condensates for autophagic processing. Disruption of this compartmentalized sorting, either at the level of scaffolds or client proteins by pathogenic mutations or stress induced proteostasis burden or autophagosome-lysosome dysfunction which alter scaffold-client interaction and enhance the retention within TRIM32 condensates, may contribute to ubiquitin-positive p62-positive inclusion formation and TDP-43 aggregation.

### TRIM32 granules accumulate in human TDP-43 proteinopathy

To further explore the disease relevance of TRIM32 condensates, we examined TRIM32 in patients with TDP-43 proteinopathy associated with different neurodegenerative diseases (Supplementary Table 1). We performed multiplex fluorescence immunohistochemistry (IF) with phospho TDP-43 (Ser409/410) to detect pathologically phosphorylated protein. We examined frontal cortex and regions of the amygdala including entorhinal cortex since these regions have been reported to exhibit early pathological TDP-43 changes in neurodegenerative diseases.^5, 76, 77^. Aged controls showed no cytoplasmic pTDP-43 aggregates and TRIM32 IF was primarily perinuclear/somatic in neurons and partially co-localized with UBQLN2 (Fig. 7A, 7B). Examination of pTDP-43 in the amygdala of six AD cases revealed that TRIM32 and UBQLN2 IF colocalized with over 80% of pTDP-43 aggregates (Fig. 7C). Colocalizing aggregates predominantly appeared as round-like cytoplasmic inclusions (Fig. 7A, 7B). In the frontal cortex, TRIM32 colocalization with pTDP-43 aggregates was less frequent (15–20%). We also observed TRIM32 co-localization with pTDP-43 inclusions lacking detectable UBQLN2 (Extended Data Fig. 6A, B). In FTLD and LATE cases with extensive pathology, diverse pTDP-43 inclusion morphologies were present. TRIM32 was not observed in association with thick neurites or skein-like inclusions (Extended Data Fig. 6C) and showed variable colocalization (15-80%) with pTDP-43 inclusions (Fig. 7C). Two relatively young cases (AD+LATE#2, 61 years; FTLD#3, 51 years) were enriched for round-like pTDP-43 inclusions that robustly colocalized with TRIM32, similar to the AD cases (Fig. 7C), suggesting TRIM32 condensates preferentially associate with round-like pathological inclusions. Consistent with in vitro reconstitution assays and cell-based observations, TRIM32 co-localized with TDP-43 inclusions and Amylo-Glo suggesting association with pathological TDP-43 amyloid (Fig. 7D).Together, these findings support a mechanistic link between TRIM32 condensates, inclusion formation, and pathological aggregation in TDP-43 proteinopathy.

## Discussion

In this study, we demonstrate that endogenous TRIM32 granules form in response to combined proteotoxic and oxidative stressors, the two major characteristics of the dual-hit hypothesis of neurodegeneration. Distinct from stress granules, which support the ISR (integrated stress response) translation programme^78^, the TRIM32-scaffolded “shuttle condensate” emerges as a regulator of proteostasis, linking normal protein quality control to neurodegenerative pathology through its stress responsiveness. Under physiological conditions, TRIM32 condensate functions as a dynamic hub, sequestering neurodegeneration-linked misfolded protein such as TDP-43 and ANXA11 and routing them toward autophagic clearance. However, this protective mechanism operates at a vulnerable kinetic boundary. When chronic stress or pathological insults impair autophagic flux, the condensate shifts from a functional transit compartment into a maladaptive trap, thereby recapitulating the multifactorial nature of TDP-43 proteinopathy.

Studies here reveal that TRIM32 functions as a scaffold for a unique “shuttle condensate”. Like other biomolecular condensates TRIM32-UBQLN2-p62 can function as a reversible sorting compartment that concentrates selected factors to coordinate trafficking and protein quality control but conspicuously engages multiple gene products linked to neurodegenerative diseases. TRIM32-UBQLN2-p62 is anticipated to traffic proteins via p62 to autophagosomes^79^ and contribute to the natural proteostasis of TDP-43, ANXA11, as well as other proteins. In pathological conditions that delay autophagy-lysosome flux^80^, condensate proteins can “age” into less dynamic states that favor aberrant aggregation when interaction energies rise or molecular exchange slows^81–83^. Findings suggest that TRIM32–UBQLN2–p62 assemblies operate at this kinetic boundary; competitive binding within the condensate governs component mobility, client residence time, and the probability of time-dependent liquid-to-solid maturation.

A defining observation is that amyloid conversion occurs in a fully reconstituted system containing only wild-type proteins under near-physiological buffer conditions, without crowding agents, agitation, or extreme ionic strength. This indicates that the core components and interaction topology are sufficient to access an amyloid-like endpoint. In cells, genetic disease(UBQLN2 mutant) and oxidative stress (arsenite) -linked perturbations accelerate this conversion. Wild-type TRIM32 with wild-type UBQLN2 forms largely spherical condensates, whereas ALS/FTLD-associated UBQLN2 variants produce enlarged, irregular assemblies that stain with Amylo-Glo, consistent with accelerated material aging in a cellular environment.

TRIM32 provides the multivalent scaffold that builds these condensates. Through E3 ligase activity, TRIM32 autoubiquitinates at multiple sites and generates mixed, branched polyubiquitin chains (K11/K48/K63). This ubiquitin topology recruits UBA domain proteins, including UBQLN2 and p62, via UBA-polyubiquitin interactions, enabling E3 activity-dependent phase separation. The scaffold creates a high-avidity compartment that can support selective capture and sorting, but it also creates the risk that prolonged residence within this environment can promote conformational sampling and nucleation events that drive liquid-to-gel/solid maturation^81–83^.

Selectivity and material state are coupled by a distinctive element in TRIM32’s substrate-binding region. A conserved hydrophobic helix/loop in the NHL domain is important for TRIM32’s normal co-functionality with UBQLN2 and for retention of UBQLN2 client proteins, including TDP-43 and ANXA11. Data support a “mimic” model in which this TRIM32 hydrophobic element competes with low-complexity interaction motifs in clients and thereby regulates client mobility through UBQLN2’s STI1 domains. This competition makes dwell time a key modifiable variable. When exchange is rapid, the compartment remains fluid and functions as a reversible sorter, but when exchange slows clients are retained longer increasing the likelihood of time-dependent amyloid conversion.

These properties define “shuttle condensates,” characterized by differential mobilities among scaffolds and clients. TRIM32’s polyubiquitin scaffold provides a selective retention environment, while UBQLN2 remains comparatively mobile and can exchange between the condensate and external pools of client proteins, enabling capture–delivery–release cycles (Fig. 7F). This architecture offers a mechanism to move UBQLN2 clients from client-rich assemblies such as stress granules into a distinct TRIM32-defined retention/sorting compartment, helping reconcile why ubiquitin-and p62-positive inclusions in patient tissue often appear outside canonical stress granules^84–87^.

ALS/FTLD-linked UBQLN2 variants disrupt this mobility-based sorting logic. Across mutant alleles tested, UBQLN2 becomes immobilized within TRIM32 condensates, condensates enlarge and become irregular, and amyloid conversion accelerates. These changes coincide with enhanced retention and aggregation of UBQLN2 clients such as TDP-43. A parsimonious explanation is that pathogenic variants, especially those in or influencing the PXX region, destabilize an intramolecular autoinhibitory conformation that normally limits STI1-I engagement^32^, freeing STI1 to bind the TRIM32 hydrophobic element with higher avidity. Increased avidity would reduce UBQLN2 exchange, collapse shuttle behavior, and bias the compartment toward pathological retention and maturation.

TRIM32’s behavior aligns with broader TRIM-family biology^54^ while providing mechanistic insight into puncta formation and proteostatic function. Many TRIM proteins form puncta and undergo autoubiquitination^54^, but TRIM32 uniquely couples strong autoubiquitination with a conserved hydrophobic element that confers selective interplay with UBQLN2 and client retention. Importantly, UBQLN2 or p62 suppress TRIM32 autoubiquitination, indicating that condensate assembly can implement negative feedback to constrain ubiquitin-chain production and stabilize scaffold architecture. Because TRIM32 builds mixed branched chains, UBQLN2/p62 may actively shape ligase output or shift chain composition and thereby influence whether cargos are routed toward proteasomal versus autophagic fates. p62’s involvement further links the compartment to autophagic delivery, making clearance efficiency a key determinant of whether sorting remains productive or evolves toward pathological maturation.

This framework predicts how diverse disease drivers converge on a shared kinetic vulnerability. Autophagy–lysosome dysfunction, as occurs in many neurodegenerative diseasesxxNixon ref, should increase residence time by impairing clearance of p62-linked cargo, promoting retention and maturation of clients such as TDP-43 and ANXA11. Stress-associated insults or factors that reduce UBQLN2 fluidity (including C9ORF72 dipeptide repeats reported to associate with UBQLN2^88^) are likewise expected to decrease exchange and accelerate maturation toward amyloid aggregates. Mutations in client proteins that alter UBQLN2-binding motifs could also shift competitive binding toward TRIM32-dependent retention, further increasing dwell time. In each case, reduced exchange would deplete functional UBQLN2 and trap clients and scaffold in immobile, aggregate-prone states, amplifying proteostatic collapse.

Human tissue observations support disease relevance and inclusion-type specificity. TRIM32 granules predominantly associate with pathologically phosphorylated TDP-43 in round-like aggregates, especially in the earliest-affected amygdala regions of patients with TDP-43 proteinopathy and in relatively young patients with FTLD and LATE, underscoring a direct link between TRIM32 condensates, inclusion formation, and pathological aggregation. Our findings provide a unifying testable pathogenic mechanism underlying UBQLN2 and TDP-43 proteinopathy across diverse neurodegenerative disorders^15, 30, 36–38^. Remarkably, our results reveal that UBQLN2 and p62 with TRIM32, traditionally viewed as clearance factors, actively promote inclusion formation and amyloid conversion, these functions far beyond expectations from prior studies^34, 40, 70, 79, 88^. This discovery challenges the current understanding of these factors and suggests that under conditions of proteostatic stress, the cell’s own quality-control machinery can be subverted into a driver of amyloid aggregation.

In sum, TRIM32-driven condensates constitute a selective sorting compartment built by E3 ligase activity, dependent polyubiquitin scaffolding, and tuned by competition between a TRIM32 mimic element, UBQLN2 STI1 mediated binding, and client low-complexity motifs. Preserved exchange supports dynamic shuttle behavior, while impaired exchange, most clearly caused by UBQLN2 pathogenic variants, shifts the system toward pathological retention, time-dependent amyloid maturation, and ubiquitin-positive inclusion formation characteristic of TDP-43 proteinopathies.

## Acknowledgments

We thank Robert O’Meally at the Mass Spectrometry and Proteomics Facility in Johns Hopkins University and Ross Tomaino at Taplin Mass Spectrometry Facility in Harvard Medical School for mass spectrometry analysis. We thank Dr. Carlos Castañeda for providing the construct UBQLN2-mCherry. Research support was provided by U19-AG065169 (P.F.W), R35NS097966 (P.F.W) and R01 MH053608 (P.F.W.). The Brain Resource Center is supported by the JHU Alzheimer’s Disease Research Center (NIH P30AG066507).

## Author contributions

W.Z. conceived and designed the project. W.Z. performed biophysical experiments and interpreted results. F.M., Z.H. and X.Y. performed plasmid extract, cell culture, human brain tissue staining and imaging. J.Z performed imaging analysis. R.Z. provided the Alphafold3 setup. D.W.N and J.C.T. provided human tissues and helped with the human brain slice analysis. W.Z., Z.H. and P.F.W wrote the manuscript with input from other authors. P.F.W and W.Z. supervised the work. All authors contributed to data analysis and interpretation.

