## Supplemental table 1 for "The TRIM32 granule promotes TDP-43 inclusion formation and amyloid aggregation through shuttle condensates"

**Supplementary Table 1. Characteristics of Controls, Alzheimer’s Disease (AD), and limbic-predominant age-related TDP-43 encephalopathy (LATE), frontotemporal lobar degeneration (FTLD) Subjects**

| **ID** | **FDX** | **CERAD** | **BRAAK** | **AGE** | **SEX** | **RACE** | **PMD** |
| --- | --- | --- | --- | --- | --- | --- | --- |
| **Control#1** | **CONTROL** | **0** | **2** | **88** | **M** | **W** | **5** |
| **Control#2** | **CONTROL** | **0** | **1** | **56** | **F** | **W** | **23** |
| **AD#1** | **AD low level, ARTAG** | **C** | **2** | **79** | **M** | **W** | **8** |
| **AD#2** | **AD intermediate level** | **C** | **3** | **>90** | **F** | **W** | **9** |
| **AD#3** | **AD high level** | **C** | **5** | **>90** | **F** | **W** | **54** |
| **AD#4** | **AD high, ARTAG** | **C** | **6** | **64** | **F** | **W** | **18** |
| **AD#5** | **AD, TDP-43 Proteinopathy** | **C** | **6** | **79** | **M** | **W** | **15** |
| **AD#6** | **AD High, TDP-43 pathology, ARTAG, CAA** | **C** | **6** | **>90** | **F** | **W** | **4** |
| **AD+LATE#1** | **AD intermediate, LATE** | **C** | **4** | **86** | **F** | **W** | **28** |
| **AD+LATE#2** | **AD intermediate, LATE 1/3, ARTAG, CAA** | **C** | **4** | **61** | **M** | **W** |  |
| **AD+LATE#3** | **AD high, LB LIMBIC, LATE 1, ARTAG** | **C** | **6** | **>90** | **M** | **W** | **28** |
| **FTLD#1** | **FTLD-TDP43(C9orf72)+motor neuron disease, AD low** | **0** | **3** | **73** | **F** | **W** | **17** |
| **FTLD#2** | **FTLD-TDP, AD prob, (no C9)** | **B** | **4** | **78** | **M** | **W** | **4** |
| **FTLD#3** | **FTLD-TDP, PART** | **0** | **1** | **51** | **M** | **W** | **38** |
| **FTLD#4** | **FTLD TDP-43, (C9orf72) CVD (NC)** | **0** | **2** | **58** | **F** | **W** | **7** |
