## Supplementary material for "The TRIM32 granule promotes TDP-43 inclusion formation and amyloid aggregation through shuttle condensates": methods

### **Construction of expression plasmids**

The DNA sequence encoding TRIM32 w/o the fluorescence protein mCeruleans3 tagged at the C-terminus was codon-optimized for expression in E. coli and inserted into expression vector pETDuet-1 by GeneUniversal. UBQLN2 with an additional cysteine at the N-terminus and His-tag at the C-terminus was constructed synthetically with codon optimization for *E. coli* expression into the vector pMAL-c5X from GeneUniversal. UBQLN2 STI1-I domain (161-297) was subcloned into vector pGEX-6P-1 (GE Healthcare) between the EcoRI and XhoI restriction sites for expression in E. coli. *pEGFP-N1_hTRIM32 (Addgene: #*69541*), HA-p62 (Addgene: #*28027*),* mCherry-p62 *(Addgene: #*187901*),* mCherry-ANXA11 (*Addgene: #*164203*),* pcDNA3.2 TDP-43 YFP (*Addgene: #*84911), pcDNA3.2 TDP-43 NLS1 YFP (*Addgene: #*84912), were purchased from Addgene. pEGFP-N1_hTRIM32 (Addgene: #69541) and UBQLN2-mCherry (from Dr. Carlos Castañeda) were used as the backbone for mCardinal-TRIM32 and UBQLN2-mCardinal, respectively. All constructs were sequenced to confirm accuracy of cloning. Bacterial plasmids obtained from Addgene include: UBE1 (Addgene plasmid # 34965), UBE2D3 (Addgene plasmid # 12643), ANXA11 (Addgene plasmid # 164496), p62 (Addgene plasmid # 190929), TDP-43 (Addgene plasmid # 133320 and plasmid # 27462). Mutations to generate substitutions of TRIM32 and UBQLN2 were introduced by NEB Q5 site-directed Mutagenesis kit (E0554S) and confirmed by the Sanger sequence or NGS (Genewiz sequence service).

### **Protein Expression and Purification**

### *E. coli* strain SoluBL21 cells (Amsbio LLC) were used for expression of all recombinant proteins. Cultures were grown in Luria-Bertani (LB) medium at 37°C to an OD_600_ (optical density at 600 nm) of 0.6 to 0.8 and induced with 0.2 mM isopropyl β-d-1-thiogalactopyranoside (IPTG) at 18°C for 12 to 16 hours. The cells were harvested and lysed by sonicating in PBS (phosphate buffered saline, pH 7.3) supplemented with 1% Triton X-100. The recombinant protein in the supernatant of the cell lysate was applied to glutathione affinity or Ni^2+^-agarose, depending on the tag system, incubated for 3 to 4 hours on a rotary shaker at 4°C and the mixture was washed extensively in PBS with 1% Triton X-100. GST-tagged proteins on glutathione affinity beads were cleaved with precision protease overnight to remove the GST tag in a buffer of 20 mM Tris-HCl (pH 8.0), 100 mM NaCl and 1 mM DTT (dithiothreitol). His-tagged proteins were eluted in 25 mM HEPES (pH 7.5), 100 mM NaCl, 5 mM β-ME, and 200 mM imidazole. Further purification was performed by size exclusion chromatography on a Superdex 75 column (GE Healthcare) and pre-equilibrated in 10 mM HEPES (pH 7.5), 100 mM NaCl, and 1 mM DTT before snap-freezing in liquid nitrogen and storage at −80°C. Protein concentrations were determined by measurement of the absorbance at 280 nm based on molar extinction coefficients calculated from the relevant sequences using Expasy’s ProtParam.

#### **GST pull-down assay**

GST fusion protein was expressed in E. coli and immobilized on glutathione-Sepharose beads. These beads were incubated with the target protein, also expressed in E. coli, in a reaction buffer at 4°C for 4 hours. Following incubation, the beads were washed three times with a buffer containing 0.5-1.0% Triton X-100 in phosphate-buffered saline (PBS) to remove non-specific binding proteins. The bound proteins were then eluted from the beads and analyzed by SDS-PAGE followed by Coomassie blue staining to visualize the interacting proteins.

### **Autoubiquitination assays**

Autoubiquitination assays were performed in a reaction buffer containing 50 mM HEPES (pH 7.5), 100 mM NaCl, 10 mM MgCl₂, and 0.5 mM DTT. Each 25 μL reaction mixture contained 40 μM ubiquitin, 100 nM E1, 2 μM E2, and 0.2 μM TRIM32, with or without the addition of 1 μM UBQLN2 and/or 0.5 μM p62 or as indicated in figure legends. The reactions were initiated by the addition of 10 mM ATP and incubated at 37°C. At various time points, reactions were quenched by adding 4x SDS-PAGE sample buffer. The samples were then resolved on 4-12% NuPAGE Bis-Tris gels. Gel bands corresponding to ubiquitinated TRIM32 were excised for mass spectrometry analysis at the Taplin Facility (Harvard Medical School) to identify ubiquitination sites on both TRIM32 and ubiquitin. All experiments were performed in at least two independent replicates with consistent results, and representative gels are shown. Gel images were acquired using a Bio-Rad Gel Doc EZ Imager with Image Lab software.

**Ub-clipping analysis of autoubiquitination reactions**

TRIM32 autoubiquitination reactions were incubated at 37 °C for 3 hour, then 10 μM Lb^pro*^ in 50 mM Tris pH 8.0, 10 mM DTT was added into TRIM32 autoubiquitination reactions for 1 hours. The reactions were diluted to 1 μM ubiquitin with quenching solution (50% acetonitrile (v/v), 0.1% formic acid (v/v)). Intact ubiquitin mass spectrometry was performed at the Mass Spectrometry and Proteomics Facility in Johns Hopkins University. Spectra were averaged and subsequently deconvoluted using Thermo Xcalibur Qual Browser (Thermo Fisher Scientific). The intensities of un- (8,451.65 Da), single- (8,565.69 Da), double- (8,679.73 Da), triple- (8,793.78 Da), and quadruple-modified (8,907.82 Da) ubiquitin were exported from Xcalibur into Microsoft Excel for further analysis. The protein samples as indicated were mixed with 4x SDS-PAGE sample buffer and resolved on NuPAGE 4–12% Bis-Tris gels and visualized by Coomassie Blue staining.

### **Phase separation assay**

Full-length recombinant ANXA11 and UBQLN2 with an N-terminal cysteine were fluorescently labeled by conjugation to Alexa Fluor 546 maleimide and Alexa Fluor 647 maleimide, respectively. Phase separation was assessed in TRIM32-mCerulean autoubiquitination reactions. The reactions were supplemented with 1 μM UBQLN2-647, 0.5 μM p62-EGFP, 0.5 μM ANXA11-546, or 0.5 μM TDP-43-mCherry, as indicated. All reactions were performed in a buffer containing 50 mM HEPES (pH 7.5) and 100 mM NaCl and were incubated for 3 hours. For imaging, 10 μL of each sample was transferred to an 18-well chambered coverslip (IBIDI). To assess amyloid aggregation within in vitro reconstituted phase separation, the amyloid cross-β-sheet dye Amylo-Glo (Biosensis, catalogue no. TR-300-AG) diluted 1:10 in 50 mM HEPES (pH 7.5) and 100mM 100 mM NaCl was mixed with 10 μl of TRIM32 auto-ubiquitination reaction in presence of 1 μM UBQLN2-647, 0.5 μM p62-EGFP, 0.5 μM ANXA11-546, and 0.5 μM TDP-43-mCherry for 10 min at indicated time points. Images were acquired using a ZEISS 880 confocal microscope equipped with a 63x oil immersion objective. Quantitative analysis of phase separation was performed using ZEN blue software.

**Fluorescence recovery after photobleaching (FRAP)**

*In vitro* FRAP was imaged on ZEISS 880 confocal with a 63X oil immersion objective. TRIM32-mCeruleans, UBQLN2-647, p62-EGFP, ANXA11-546 or TDP-43-mCherry were imaged before and directly after bleaching with a laser intensity of 100% at 405 nm (for mCeruleans), 480 nm (for EGFP), 530 nm (for ANXA11-546 or mCherry) or 647nm (for UBQLN2-647). And the fluorescence recovery of the bleached region was measured as indicated time. For bleached region of interest (ROI), a background ROI and a non-bleached droplets were recorded in parallel.

### **Droplet sedimentation assay**

To determine the accumulation of ubiquitinated protein within condensates, we performed an in vitro sedimentation assay. Samples from 3-hour TRIM32 autoubiquitination reactions in the presence of 1 μM full-length recombinant UBQLN2 or 0.5 μM p62 were centrifuged at 12,000 g for 5 minutes at 4°C. The supernatant and pellet fractions were separated. The pellet was then washed once with HEPES buffer to remove non-condensed proteins. Both the supernatant and pellet fractions were analyzed by SDS-PAGE followed by western blotting. Ubiquitinated proteins were detected using an anti-ubiquitin antibody (Santa Cruz Biotechnology: sc-8017).

**Western blotting**

Samples were resolved by SDS-PAGE using a 4–12% NuPAGE Bis-Tris Gel alongside a protein ladder and running with constant voltage of 120 V. Proteins were then transferred onto a PVDF membrane at a constant current of 450 mA for 2 hours. The membrane was blocked for 1 hour at room temperature with 5% nonfat dry milk (Biorad-1706404) in TBST buffer (1x TBS, Biorad, 161-0782 with 0.1% Tween 20). Subsequently, the membrane was incubated overnight at 4°C with a primary antibody. Primary antibodies used were: Ubiquitin (Santa Cruz, catalogue no: sc-8017), TDP-43 (Abcam, catalogue no: Ab104223), Annexin11 (Proteintech, catalogue no: 10479-2-AP). Following a series of washes, the membrane was incubated with the appropriate secondary antibody (1:1000 dilution in TBST) for 1 hour at room temperature. A loading control antibody was also used for normalization. Protein bands were visualized using chemiluminescence and imaged on a Bio-Rad ChemiDoc MP Imaging System.

**Co-Immunoprecipitation (CoIP) Assay**

Brain tissue from 6-month-old C57BL/6 mice was harvested and homogenized in an immunoprecipitation (IP) buffer containing 1% Triton X-100 in PBS, supplemented with PhosSTOP phosphatase inhibitor and a protease inhibitor cocktail (Roche, Switzerland). The lysates were then sonicated with 10 pulses of 0.4 seconds each and centrifuged at 12,000 g for 10 minutes to remove insoluble material. The resulting supernatant was incubated with the indicated primary antibodies and Dynabeads Protein G overnight at 4 °C with gentle rotation. Following incubation, the beads were washed three times with the IP buffer to remove non-specific binding proteins. Proteins were then eluted from the beads by boiling in 2x SDS-PAGE sample buffer and analyzed by SDS-PAGE and subsequent western blotting. Primary antibodies used for IP: UBQLN2 (Santa Cruz, sc-100612).

**Cell culture, transfection, and immunofluorescence**

HEK293 cells were cultured in Dulbecco’s modified Eagle’s medium (DMEM, Life Technologies) supplemented with 10% fetal bovine serum (FBS) and 50 µg/mL penicillin-streptomycin. Cells were maintained at 37°C in a humidified incubator with 5% CO₂. Transfections were performed using X-tremeGENE™ HP DNA Transfection Reagent (Sigma) according to the manufacturer's protocol. To induce stress, cells were incubated in complete media supplemented with 0.2 mM sodium arsenite (Titripur, 1062771003; stored as 50 mM aqueous stock) for 120 min at 37 °C.

For immunofluorescence analysis, cells were fixed with 4% paraformaldehyde (PFA) in PBS for 15 minutes at room temperature (RT). Cells were then permeabilized with 0.2% Triton X-100 in PBS for 10 minutes at RT, followed by blocking with 10% normal goat serum for 1 hour at RT. After blocking, cells were incubated with the indicated primary antibodies overnight at 4°C. Following three washes (5 minutes each) with PBS, cells were incubated with a fluorescently labeled secondary antibody (1:1000 dilution in PBS) for 1 hour at RT. Cells were then washed twice with PBS before being stained with DAPI for 10 minutes at RT. For TDP-43 amyloid fibril staining, cells were fixed with PFA, then transferred into 70% ethanol for 5 min at RT. After rinsed in distilled water for 2 min, the slides were incubated with Amylo-Glo (Biosensis, catalogue no. TR-300-AG, 1:100) for 10 min, followed by two washes with saline; cells were not stained with 4′,6-diamidino-2-phenylindole. Coverslips were mounted using Prolong Gold antifade reagent (Invitrogen #P10144) and stored at 4°C in the dark until imaging.

Images were acquired with ZEISS LMS 880 or ZEISS LMS 800 confocal microscope image and analyzed using FIJI/ImageJ software. Primary antibodies: UBQLN2 (Santa Cruz, sc-100612).

**Differential solubility assay**

Transfected HEK293T were harvested on ice using cold RIPA buffer with PhosSTOP phosphatase inhibitor and a protease inhibitor cocktail (Roche, Switzerland). Following lysis on ice for 15 min with occasional inversion, Cells were sonicated at 30% amplitude for 0.4 s 8x. Samples were centrifuged at 28,000g for 15 min at 4°C to pellet RIPA-insoluble material, with the supernatant removed and saved as the RIPA-soluble fraction. The RIPA-insoluble pelletswere washed twice with RIPA buffer, with centrifugation at 28,000 g for 15 min at 4 °C after each wash. Pellets left were resuspended in urea buffer (8 M urea, 10 mM Tris-HCl, 100 mM sodium dihydrogen phosphate, pH 8.0) as the RIPA-insoluble, urea-soluble fraction.

**Primary mouse cortical neuron culture and immunofluorescence**

Cortexes were dissected from embryonic day 18 (E18) C57/Bl6 mouse pups and dissociated with papain and DNase. Cortical neurons were plated at a density of 3 × 10^5^ cells per well on 18 mm cover slips coated with poly-L-lysine in 12-well cell culture plates. On DIV (day in vitro.)1-3, neurons were maintainedin Neurobasal medium (Thermo Fisher Scientific) containing 5% FBS, 1% penicillin/streptomycin (Thermo Fisher Scientific), 2% Glutamax-I (Thermo Fisher Scientific) and 2% B-27 supplement (Thermo Fisher Scientific). Cytosine arabinoside (AraC) was added on DIV 4 to inhibit glial proliferation, and half of the medium was replaced with glial conditional Neurobasal medium containing 1% FBS, 1% penicillin/streptomycin, 2% Glutamax-I and 2% B-27 supplement. Thereafter, half of the medium was replaced every 3 days. On DIV 14, neurons were treated with sodium arsenite (200 μM for 4 h), bicuculline (40 μM for 4 h or 24 h), and/or bafilomycin A1 (10 nM for 4 h), as indicated in the relevant figure panels.

For immunofluorescence, neurons were fixed in 4% paraformaldehyde in PBSfor 10 min, permeabilized with 0.2% Triton X-100 in PBS for 10 min, and blocked with 5% bovine serum albumin (BSA) for 1 h. Primary antibodies were diluted in 5% BSA and incubated with neurons overnight at 4°C. Fluorescence dye-conjugated secondary antibodies to the appropriate species were diluted in FBS and incubated at room temperature for 1 hr. Nuclei were counterstained with DAPI. Primary antibodies include UBQLN2 (Santa Cruz, sc-100612, 1:400), pTDP-43 (Ser409/410) (Biolegend, BL829901, 1:400), TRIM32 (Invitrogen, PA5-22316, 1:500).

**Human tissue samples and immunofluorescence staining**

All human tissue samples were obtained from the Johns Hopkins Brain Resource Center in Baltimore, MD, with a post-mortem interval of less than 24 hours (see supplement table 1 for details). Control cases were selected from individuals who were matched for age to AD and FTD subjects and showed no evidence of amyloid deposition or phosphorylated TDP43. AD and FTD cases were chosen based on evidence of TDP-43 pathological level and were also age-matched to the control cases.

To prepare formalin-fixed, paraffin-embedded tissue sections for analysis, 10 µm sections of the frontal cortex were cut and placed in an oven at (60°C) for 30 min. The sections were then deparaffinized into four changes of xylene for 2 minutes each, followed by rehydration in four changes of 100% alcohol and four changes of 95% alcohol for 2 minutes each. After rinsing in tap water for 5 minutes, the sections were treated with LED system for 16 - 18 hours to quench autofluorescence^1^. To retrieve antigens, the sections were incubated in Antigen retrieval solution (10 mM sodium citrate buffer with 0.05% Tween 20, pH 6.0) in 95°C water bath for 30 min and cooled down to RT for 100 min. After antigen retrieval, the sections were washed in 1X PBS for 5 minutes for three times. Wax borders were drawn with an ImmEdge Hydrophobic Barrier PAP pen. The sections were permeabilized in PBS-T (PBS with 1% TrionX-100) for 15 min at 4°C and then blocked with a blocking buffer containing 5% normal goat serum and 0.3% Triton X-100 in PBS at room temperature for one hour. For Amylo Glo staining, the slides are transferred into a 70% solution of ethanol for 5 minutes at room temperature. The slides are then rinsed in distilled water for 2 minutes and then incubated for 10 minutes in prepared 1X staining solution (1:100 in saline). After washing in saline solution for 5 minutes, primary antibodies were incubated overnight at 4°C, and the sections were subsequently washed three times with PBS for 5 minutes. After washing, the sections were incubated with a secondary antibody for one hour at room temperature and washed again three times with PBS for five minutes. To further quench lipofuscin, biotium Trueblack (prod.#23007) at a dilution of 1:20 was applied with a 70% alcohol solution (200 µL/slide) and swirled by hand for 50 seconds. The sections were then washed three times in PBS for 10 minutes, on a shaker plate. The slides were coverslipped using 50 µL Prolong Gold Antifade with or without DAPI. Images were taken at 40X or 63X magnification on a ZEISS LSM 880 or 800 confocal microscope, with 4-6 images taken per slice. Tiled images were taken at 40X magnification for quantification. Each staining procedure included samples processed without primary antibody and only with secondary antibodies to serve as a negative control. Image acquisition parameters were uniform across the entire cohort, and ImageJ software was utilized to quantify immunoreactive signals from images, with background secondary antibody-only signals subtracted. Brightness, contrast, and threshold adjustments were applied in the same manner across all comparison images, including the secondary-only controls. Experimenters were blinded during imaging and processing.

The following antibodies were used to probe target proteins: UBQLN2 (Santa Cruz, sc-100612, 1:200), pTDP-43 (Ser409/410) (Biolegend, BL829901, 1:200), TRIM32 (Invitrogen, PA5-22316, 1:400).

**Quantification and statistical analysis**

No statistical methods were used to predetermine the sample sizes. No data was excluded from the analysis. No formal randomization techniques were used. Samples were allocated randomly to experimental groups and processed in an arbitrary order. Means ± s.e.m. were used to represent all continuous values, as indicated in the figure legends. Statistical information for each experiment, including the statistical methods, the P values and numbers are shown in the figures and corresponding legends. Statistical analyses were performed in GraphPad Prism v.9 on data acquired from at least three independent experiments. Grouped samples were compared using One-way ANOVA followed by Tukey’s multiple comparisons test. Matched samples were compared using two-tailed Student’s paired t-test, in which n.s. (p-value>0.05), ∗ (0.01<p-value≤ 0.05), ∗∗ (0.001<p-value≤ 0.01) and ∗∗∗ (p-value≤ 0.001).
