## Supplemental figures for "The TRIM32 granule promotes TDP-43 inclusion formation and amyloid aggregation through shuttle condensates"

### Extended Data Figure 1

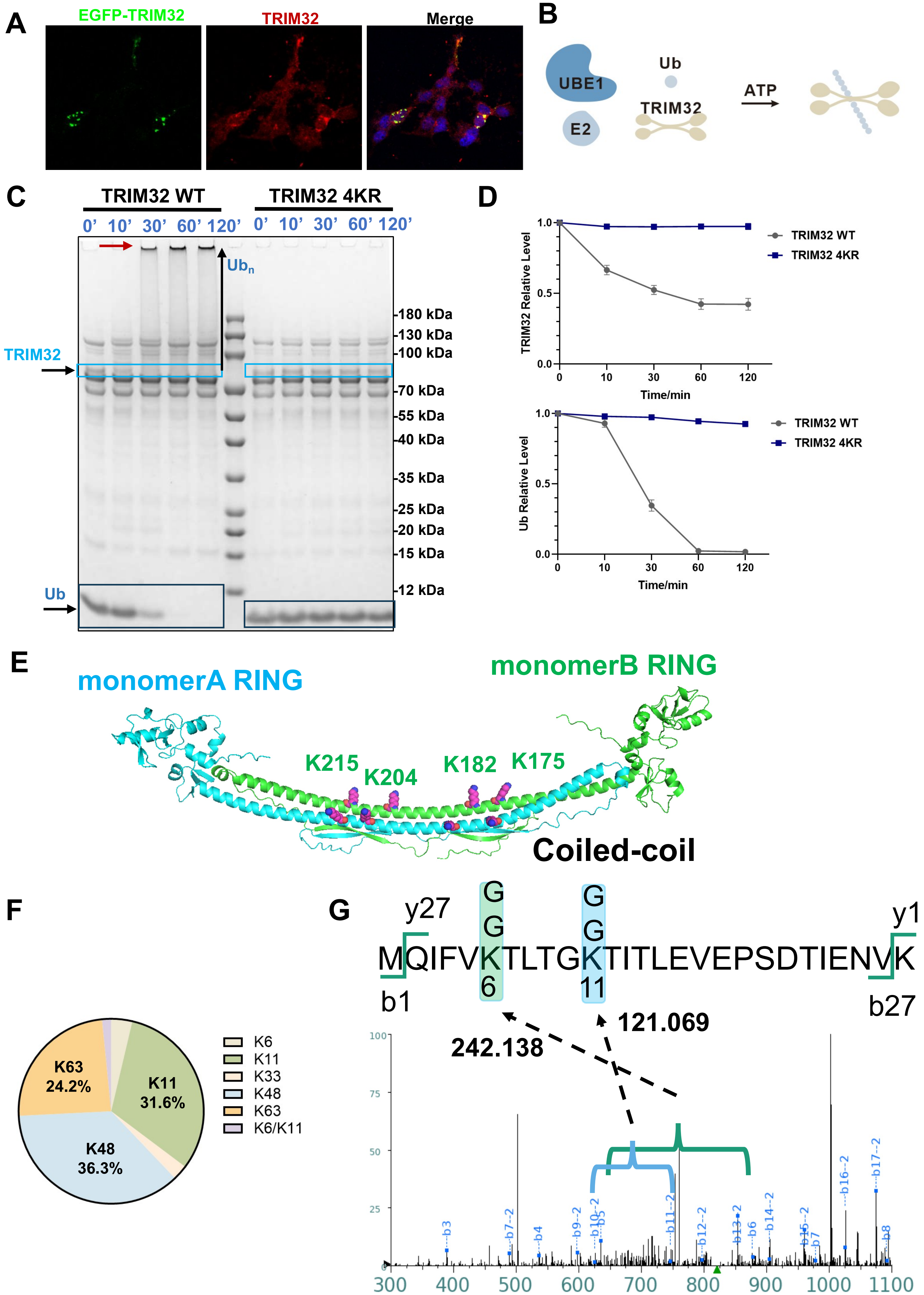

**Extended Data Figure 1. Characterization of TRIM32 assembly ubiquitin chain.**

**(A)** Validation of TRIM32 antibody for immunofluorescent staining. HEK293T cells overexpressing EGFP-TRIM32 were stained with anti-TRIM32 antibody (red).

(**B**) Depiction of the core components of the TRIM32 in vitro auto-ubiquitination reconstitution assay including UBE1, the E2 enzyme UBE2D3, ubiquitin, TRIM32, and ATP.

**(C)** Coomassie stained SDS-PAGE gels of *in vitro* TRIM32 autoubiquitination assays showing that compared with WT TRIM32, the quadruple point mutation TRIM32 4KR (K175R/K182R/K204R/K215R) prevents autoubiquitination. Colored box indicates the band for TRIM32 (cyan) and ubiquitin (blue) level quantification, respectively. Red arrow indicates HWM polyubiquitinated TRIM32 in the loading well.

**(D)** Quantification of time course change in (C) measuring TRIM32 or ubiquitin level via densitometry. Combined data from three independent experiments are shown. Data are presented as mean ± s.e.m.

**(E)** Four lysine residues (K175, K182, K204, K215) in purple identified as autoubiquitination sites cluster on TRIM32 3D structure of dimerized coiled-coil domain predicted by Alphafold.

**(F)** Ubiquitin linkage composition by AQUA MS from *in vitro* TRIM32 autoubiquitination assays performed in technical duplicate. .

**(G)** Representative MS/MS spectra of peptides from autoubiquitinated TRIM32 demonstrating ubiquitination modification concurrent at lysine 6 and lysine 11 of the ubiquitin. Peaks matching expected singly charged (+) b and doubly charged (2 +) b ions are labeled. The mass difference between the b5 and b6 ions corresponds to K6-GG. The mass difference between the b11-2 and b10-2 ions indicates K11-GG.

### Extended Data Figure 2

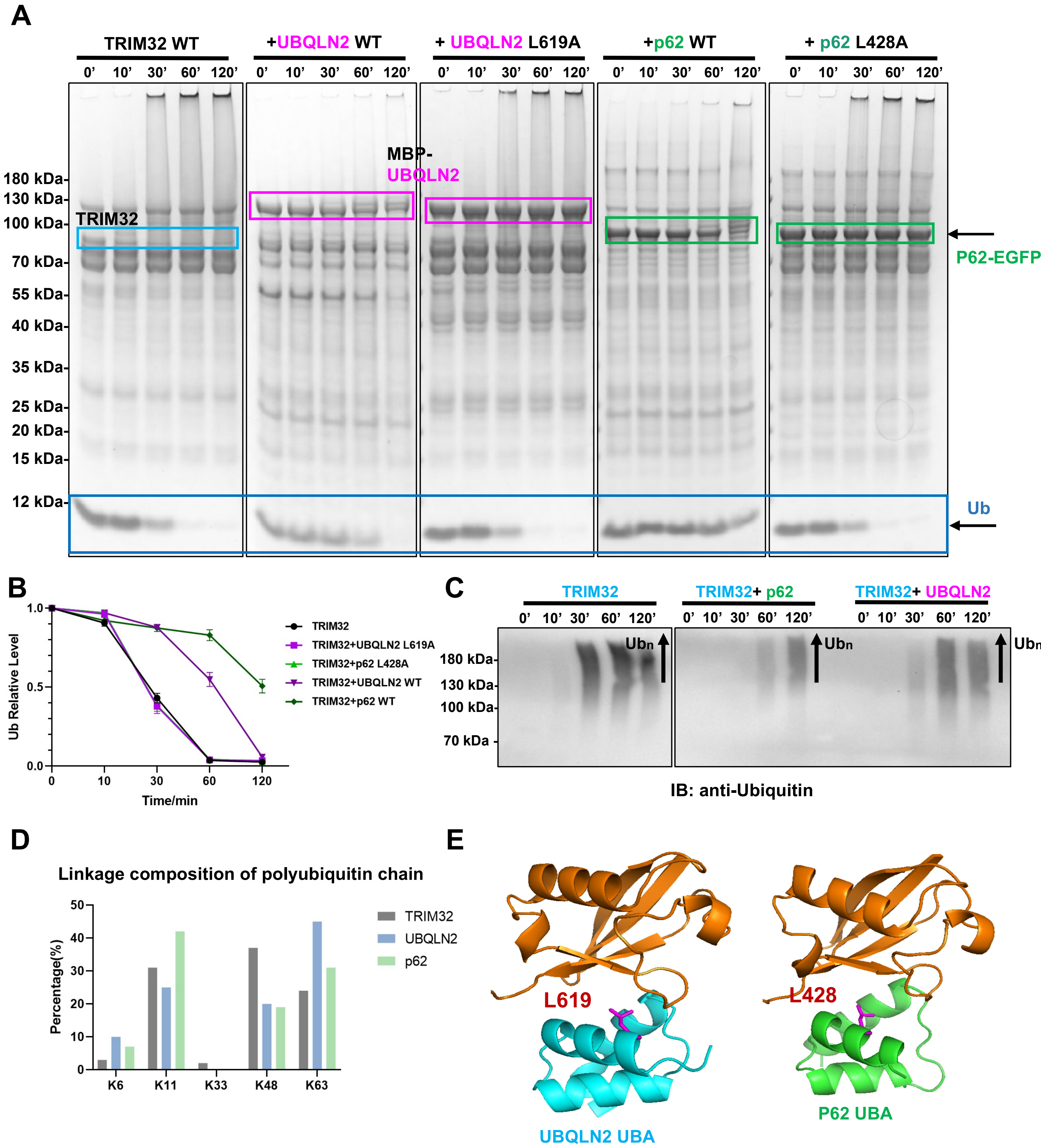

**Extended Data Figure 2. TRIM32 E3 ubiquitin ligase activity modulated by UBQLN2 and p62**

**(A)** Coomassie stained SDS-PAGE gels of *in vitro* TRIM32 autoubiquitination assays showing that the presence of WT UBQLN2 or WT p62 or their UBA domain point mutations impacts TRIM32 ligase activity, indicated by the remaining ubiquitin over the timing course. Colored box indicates the band for UBQLN2 or p62 and ubiquitin level quantification, respectively.
**(B)** The graph shows quantification of time course in **(A)** measuring ubiquitin via densitometry. Shown are combined data from three independent experiments. Data are presented as mean ± s.e.m.
**(C)** TRIM32 autoubiquitination reaction with or without UBQLN2 or p62 at indicated time assayed by ubiquitin western blot showing that UBQLN2 or p62 restrain TRIM32 mediated ployUbiquitin chain (Ubn) formation.
**(D)** Relative abundance of Linkage composition of polyubiquitin chain formed by TRIM32 with/without the addition of UBQLN2 and p62.
**(E)** Magnified view of conservative interaction interfaces between ubqln2 or p62 UBA domains and ubiquitin generated by Alphafold. The residues L619A at UBQLN2 UBA domain and L428 at p62 domain important for interaction are shown as a stick representation in purple.

### Extended Data Figure 3

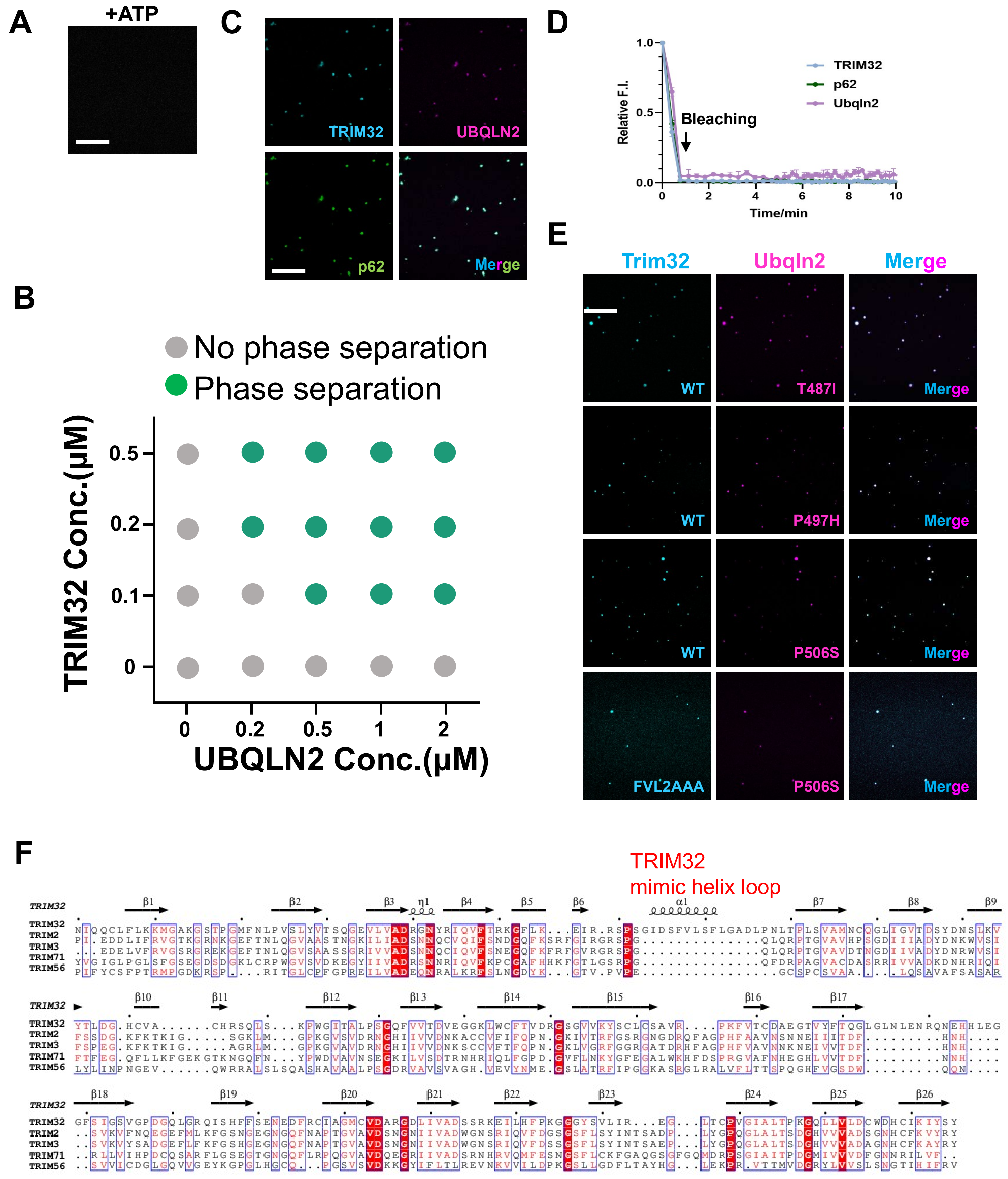

**Extended Data Figure 3. Characterization of TRIM32 ligase activity dependent phase separation.**
**(A)**TRIM32 alone cannot form phase separation in TRIM32 in vitro autoubiquitination reaction. Scale bar, 20 μm.
**(B)** Phase diagram of the formation of TRIM32 condensates at indicated TRIM32 and UBQLN2 concentrations.
**(C)** TRIM32 ligase activity-dependent formation of phase separation in TRIM32 in vitro autoubiquitination reaction in presence of UBQLN2 and p62. Scale bar, 20 μm.
**(D)** Quantification of fluorescence intensity recovery of photobleached TRIM32, p62 and UBQLN2 in **(C)**. $n = 3$. Data are presented as mean ± s.e.m.
**(E)** Formation of phase separation in TRIM32 in vitro autoubiquitination reaction with the addition of UBQLN2 pathogenic mutants or using TRIM32 mutant FVL2AAA (F417A/V418A/L419A) at the TRIM32 hydrophobic loop in the hydrophobic loop. Scale bar, 20 μm.
**(F)** The sequence alignment of the NHL domains of human TRIM family members including TRIM32, TRIM2, TRIM3, TRIM56 and TRIM71, showing the unique hydrophobic helical loop only present in TRIM32. Secondary structures of TRIM32 NHL domain are shown on the top.

### Extended Data Figure 4

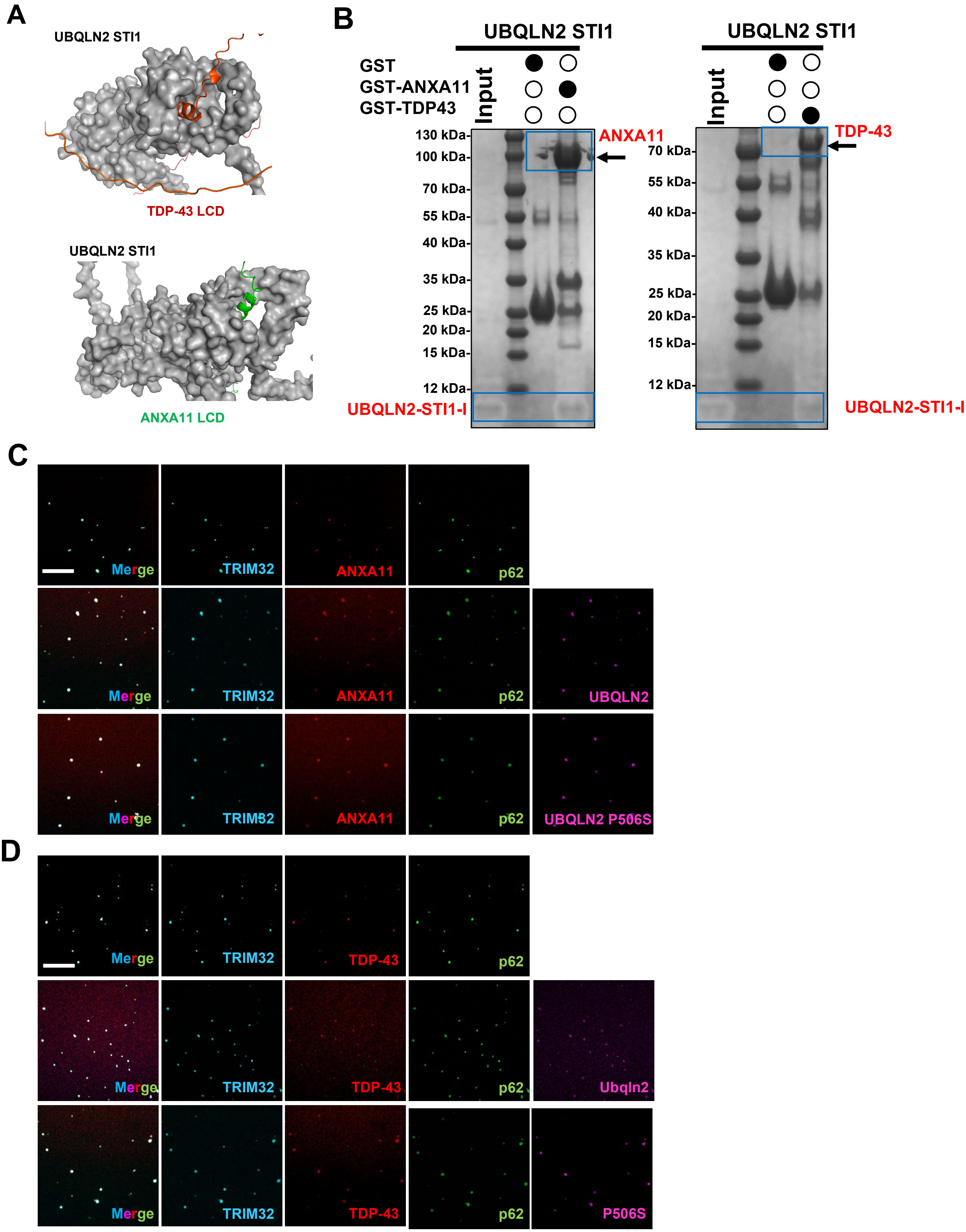

**Extended Data Figure 4. TRIM32 mimics the α-helix region of Client proteins ANXA11 and TDP-43**

**(A)** Complex structures of UBQLN2 STI1 domain and ANXA11 LCD or TDP-43 LCD generated by AlphaFold, shows the helix in red or green may occupy the same binding pocket on UBQLN2 STI1-I domain.

**(B)** Coomassie stained SDS-PAGE gels of in vitro GST pull-down showing ANXA11 or TDP-43 binds UBQLN2 STI1-I domain, which stained less intensely due to its acidic isoelectric point and small size.

**(C, D)** Formation of phase separation in TRIM32 in vitro autoubiquitination reaction in presence of p62 and ANXA11 **(C)** or TDP-43 **(D)** with the addition of wt UBQLN2 or UBQLN2 pathogenic mutant P506S. Scale bar, 20 μm.

### Extended Data Figure 5

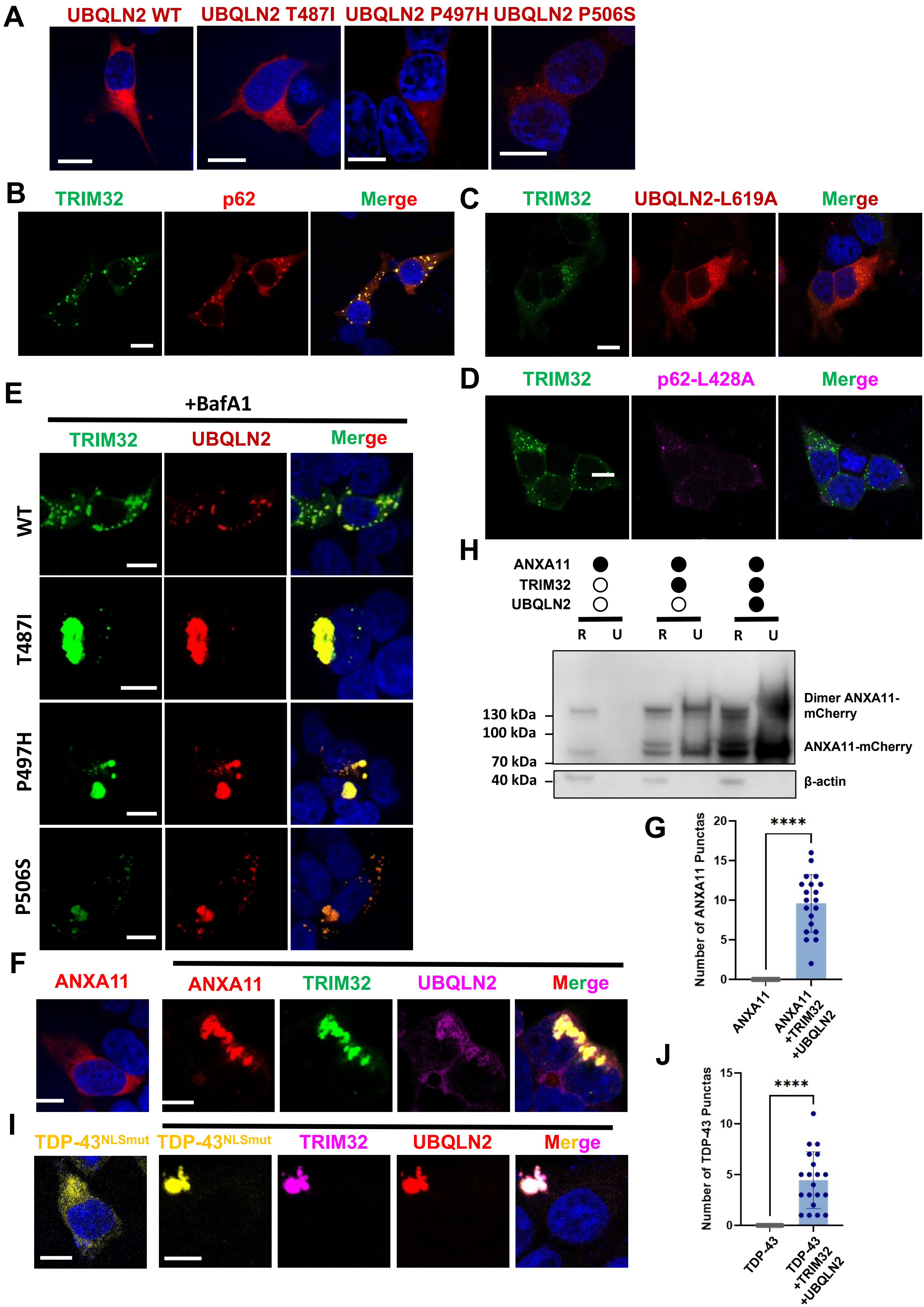

**Extended Data Figure 5. Characterization of TRIM32 dependent condensates.**

**(A)** Representative immunofluorescence images of HEK293 cell that transfected with UBQLN2 or UBQLN2 pathogenic mutant. Scale bar, 10 μm.

**(B)** Representative immunofluorescence images of TRIM32 (green) and p62 (Red) with DAPI (blue) in HEK293T cells. Scale bar, 10 μm.

**(C, D)** Representative immunofluorescence images of TRIM32 (green) and UBQLN2 UBA domain mutation L619A **(C)** or p62 UBA domain mutation L428A **(D)** with DAPI (blue) in HEK293T cells. Scale bar, 10 μm.

**(E)** Representative immunofluorescence images of TRIM32 (green) and wt UBQLN2 or UBQLN2 pathogenic mutants (Red) with DAPI (blue) in HEK293T cells. The cells were treated with Bafilomycin A1 (10nM, 24 hours). Scale bar, 10 μm.

**(F)** Representative immunofluorescence images of HEK293T cells with alone transfection of ANXA11 (red) or cotransfection of ANXA11 (red), TRIM32 (green) and UBQLN2 (Purple) with DAPI (blue). Scale bar, 10 μm.

**(G)** Quantification of number of ANXA11 puncta colocalization with TRIM32-UBQLN2 condensates showed in **(F)**.

**(H)** Representative western blot images to assess the solubility of ANXA11 with or without cotransfection of TRIM32 or UBQLN2 in RIPA (R) and urea (u) buffers, using ANXA11 antibody.

**(I)** Representative immunofluorescence images of HEK293T cells with alone transfection of TDP-$43^{NLSmut}$ (yellow), or cotransfection of TDP-$43^{NLSmut}$ (yellow), TRIM32 (purple) and UBQLN2 (red) with DAPI (blue) in HEK293T cells. Scale bar, 10 μm.

**(J)** Quantification of number of TDP-43 puncta colocalization with TRIM32-UBQLN2 condensates showed in **(I).**

Data are presented as mean ± s.e.m. Statistical significance was determined using pairwise comparisons using unpaired two-tailed t-tests. $p < 0.05$ (*), $p < 0.01$ (**), $p < 0.001$ (***), $p < 0.0001$ (****); ns, not significant. Each spot represents puncta number in one cell. Shown are combined data from three independent experiments.

### Extended Data Figure 6

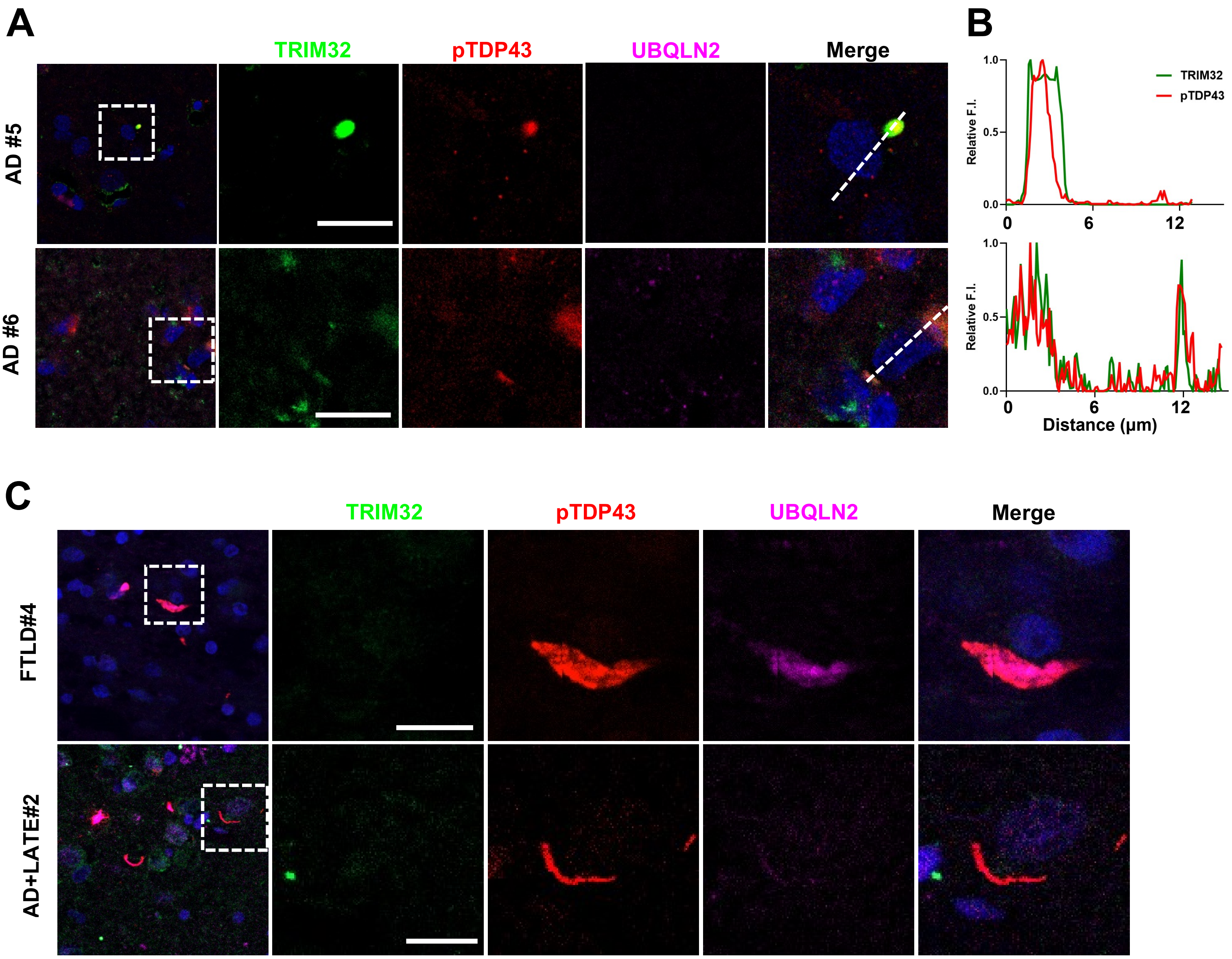


**Extended Data Figure 6. Diversity of TRIM32 condensates in patients with TDP-43 proteinopathy.**

**(A)** Representative images of TRIM32 and pTDP-43(Ser409/410) in human AD frontal cortex with TDP-43 proteinopathy. Areas outlined by dotted white boxes are magnified to the right. Scale bar, 10 μm.

**(B)** Radial distribution of the fluorescence intensities of TRIM32 and pTDP-43 indicated that TRIM32 co-localization with pTDP-43.

**(C)** Representative images showing TRIM32 and UBQLN2 within pTDP-43 thick neurites or skein-like inclusions in human C9orf72 linked FTLD with TDP-43 proteinopathy, and AD with LATE. Areas outlined by dotted white boxes are magnified to the right. Scale bar, 10 μm.
